# Dynamic Interplay of Autophagy and Membrane Repair During *Mycobacterium tuberculosis* Infection

**DOI:** 10.1101/2022.12.19.521111

**Authors:** Jacques Augenstreich, Anna T. Phan, Charles N.S. Allen, Anushka Poddar, Lalitha Srinivasan, Volker Briken

## Abstract

Autophagy plays a crucial role in the host response to Mycobacterium tuberculosis (Mtb) infection, yet the dynamics and regulation of autophagy induction on mycobacterial phagosomes remain partially understood. In this study, we employed time-lapse confocal microscopy to investigate in real time the recruitment of LC3B (LC3), a key autophagy marker, to Mtb-containing vacuoles (MCVs) at the single cell level with our newly developed workflow for single cell and single MCV tracking and fluorescence quantification. The results reveal that approximately 70% of MCVs exhibited LC3 recruitment but was lost in about 40% of those MCVs. The LC3 recruitment to MCVs displayed a high variability in timing that was independent of the size of the MCV or the bacterial burden. Most notably, the LC3-positive MCVs did not acidify, indicating that LC3 recruitment does not necessarily lead to the formation of mature autophagolysosomes. In addition, interferon-gamma (IFN-γ) pre-treatment did not affect LC3 recruitment frequency or autophagosome maturation, but increased the susceptibility of the macrophage to Mtb-induced cell death. Instead, LC3 recruitment and lysotracker staining were mutually exclusive events alternating on some MCVs multiple times showing a new reversible aspect of this autophagy response. It also suggested a role of autophagy in membrane repair of the MCV. Consistently, LC3 recruitment was strongly associated with galectin-3 and oxysterol-binding protein 1 staining, indicating a correlation with membrane damage and repair mechanisms. However, knockdown of ATG7 did not impact membrane repair, suggesting that autophagy is not directly involved in this process but is coregulated by the membrane damage of MCVs.

In summary, our findings provide novel insights into the dynamic and variable nature of LC3 recruitment and autophagy to the MCVs over time during Mtb infection. Our data suggests that there is no major role of autophagy in cell autonomous defense against Mtb nor membrane repair of the MCV in human macrophages. However, the combined dynamics of LC3 recruitment and Lysoview staining emerged as promising markers for future research focused on directly investigating the damage and repair processes of phagosomal membranes.

## Introduction

*Mycobacterium tuberculosis* (Mtb) is the causative agent of the pulmonary disease tuberculosis, which is one of the deadliest bacterial infections worldwide (WHO). One of the main niche of replication for Mtb during infection is the macrophage.^1–4^ An important characteristic of intracellular Mtb is its ability to evade host cell microbicidal responses.^5–7^ One mechanism of the cell-intrinsic host defense is the targeting of the phagocytosed bacteria by the host cell autophagy to create an autophagosome through a process called xenophagy.^8^ An important marker for autophagy activation is the protein microtubule-associated proteins 1A/1B light chain 3B (LC3), which is recruited to the membrane of the forming phagophore for the final formation of the autophagosome.^8^ The recruitment of LC3 to the Mtb phagosomes can also happen independently of the canonical autophagic machinery in a process called LC3-associated phagocytosis (LAP).^9^ There is evidence that xenophagy limits Mtb proliferation in murine and human macrophages.^10–12^ Nevertheless, the role of xenophagy in cell autonomous defense against Mtb during *in vivo* infections is more complex with some reports showing a more indirect role of autophagy for in vivo host resistance.^13, 14^ LAP also has a protective role in murine macrophages^15^, but its capacity to inhibit Mtb replication in human macrophages has not been clearly demonstrated.

Even though they both require autophagy machinery, xenophagy and LAP are triggered in distinct ways. During Mtb infections, xenophagy is mainly induced by membrane damage to the Mtb-containing vacuole (MCV), which allows the recruitment of galectin proteins that recruit proteins of the autophagic machinery^11, 16–19^. In contrast, LAP is independent of membrane damage induction and relies on TLR signaling^20^, and LC3 conjugation to the phagosome is dependent on the production of reactive oxygen species (ROS) by the NOX2 complex^21^. Ultimately, both LAP and xenophagy activation is followed by phagosome/autophagosome maturation^9^. The maturation is mainly reflected by acidification of the autophagosome which is required for creating a bactericidal microenvironment and the activity of the lysosomal proteases and hydrolases.^22^ This maturation occurs even though LC3 and autophagic machinery proteins were found to be dispensable for phagosome maturation occurring independently of xenophagy as characterized by acidification or LAMP1 recruitment.^23^

Mtb infection leads to the activation of autophagy signaling, but bacteria can inhibit steps of xenophagy and LAP to increase their survival within macrophages.^6, 7^ In the model system of *Dictyostelium discoideum* infected by *Mycobacterium marinum* (Mma), the phagosome escape by Mma triggers xenophagy, but the maturation of the autophagosome is blocked by the bacteria.^24, 25^ This inhibition of the maturation of the autophagosome was also observed in Mtb-infected human dendritic cells^26^ and macrophages^27^ and is, at least partially, mediated by the Mtb cell envelope lipid phthiocerol dimycocerosate (DIM/PDIM).^28^ The Eis and PE-PGRS47 proteins are among several Mtb proteins that have been identified to mediate the inhibition of host cell xenophagy (for review^6^).^29–32^ The observation of xenophagy via electron microscopy highlighted that Mtb could escape from the newly formed autophagosome.^33^ Mtb inhibited LAP via CpsA; however, the inhibition of LC3 recruitment was rescued by priming the macrophages with interferon-Gamma (IFN-γ).^15^

Mtb manipulates many host cell signaling pathways in order to survive and replicate in macrophages.^6, 7^ A hallmark discovery was the observation that Mtb inhibits the maturation process of the MCV.^34^ Many additional reports established a dogma that Mtb remains within an immature phagosome during its intracellular infection phase.^35–39^ This dogma was overturned when it was shown that a fraction of the bacteria can escape the MCV into the cytosol.^40, 41^ Additional studies show that a fraction of MCVs do not inhibit phagosome maturation but acidify and obtain characteristics of phagolysosomes.^40, 42^ It is not uncommon for intracellular bacterial pathogens to occupy several intracellular niches.^43^ What is the interaction of these different populations of bacteria within the infected cell and how does the niche that Mtb occupies determine its interaction with the host cell autophagy machinery? For example, the contact of the bacteria with the cytosol triggers ubiquitination of Mtb involving the E3 ligase Parkin and the DNA sensor STING, that can trigger xenophagy in an attempt to recapture the bacteria.^11, 18^ Autophagy induction is proposed to directly control Mtb infection in macrophages by increasing the killing of the bacteria in an mature autophagolysosome. However, with Mtb able to escape the autophagosome, and the mounting evidence that *in vivo*, the role of autophagy might be indirect via some immune system modulation, the question remains of how directly bactericidal autophagy is. To start answering these questions we performed a spatiotemporal analysis of the interaction of individual bacteria with the host cell LC3 protein as a marker for autophagy responses.

The crosstalk between Mtb and autophagy (*i.e* xenophagy, LAP) is highly dynamic and diverse, showing a duality between induction and inhibition of these responses by the bacteria. Live cell imaging coupled with time-lapse acquisition at the single-cell level is an excellent approach to investigate host pathogen interactions in time and space^44^ but has only recently been applied to study Mtb infections.^3, 12, 33, 45–47^ In this study, we monitored the dynamics of LC3 association to the phagosome during the infection of THP-1 cells with Mtb using time lapse confocal microscopy. We also implemented our recently release workflow combining different methods of deep learning-based segmentation and rational tracking of cells to analyze fluorescence signals at the single-cell level and/or at the single bacteria level.^48^ The results show that LC3 recruitment to the MCV is frequent but in many cases only temporary. We demonstrate that bacterial burden does not influence LC3 recruitment but may accelerate its occurrence in the cells showing the recruitment. Next, we observed that the priming of cells with IFNγ did not affect LC3 recruitment frequency nor acidification of the MCV. However, in most cases LC3 recruitment was preceded by a loss in acidification of the MCV followed by a restoration of acidity when LC3 is recruited suggesting a role of autophagy in membrane repair of the MCV. Nevertheless, knock-down approaches showed that LC3 recruitment is actually not involved in membrane repair but is concomitantly activated with the endoplasmatic reticulum (ER) dependent membrane repair machinery.

## Results

### The recruitment of LC3 is frequently observed around the MCVs over the course of infection

Multiple studies have shown that Mtb can induce autophagy at specific time points post-infection. Autophagy was evaluated by immunofluorescence staining on fixed samples^17, 28^ or stable expressions of recombinant fluorescence proteins,^33, 45^ but for the latter lacked clear quantification and timing determination. Consequently, these studies offered an incomplete view of the dynamic events and, for example, did not allow to determine if LC3 recruitment to the MCV follows a specific timing after phagocytosis or if it is a transient phenomenon.

To evaluate the dynamics of autophagy induction during infection with Mtb, we used PMA-differentiated THP-1 cells stably expressing LC3-GFP^49^ and infected them with a fluorescent strain of Mtb (H37Rv-DsRed). The cells were followed by time-lapse confocal microscopy during 16-20 hours after the addition of bacteria at on MOI of 2 (Figure 1A, Movie S1). The LC3 recruitment to the MCV was monitored from the time of phagocytosis to the end of recording for individual MCV. The results showed that ∼70% of MCVs triggered LC3 recruitment (Figure 1B) at one point or another after phagocytosis. However, almost half of the LC3-positive (LC3^+^) MCVs became negative by the end of the recording showing efficient escape of Mtb from the LC3 recruitment. We also observed two patterns of escape from LC3 recruitment by Mtb (Figure 1C-D). One pattern (Exclusion) which is characterized by an initial increase of the LC3 signal at the MCV that quickly diffused away from the bacteria (Figure 1C, Movie S2). The other pattern observed (Egress) is characterized by the accumulation of LC3 signal at the MCV with the formation of tubulovesicular structures (TVS),^50^ then the bacteria and the TVS separate (Figure 1D, Movie S3). An example of continuous recruitment of LC3 to the MCV is shown with the formation of an Mtb-containing autophagosome (Figure 1E, Movie S4). Overall, LC3 recruitment seemed frequent around MCVs (∼70% of all observed MCVs), but in a significant fraction of these recruitments (about 40%) the bacteria appear to escape from the response.

**Figure 1:**
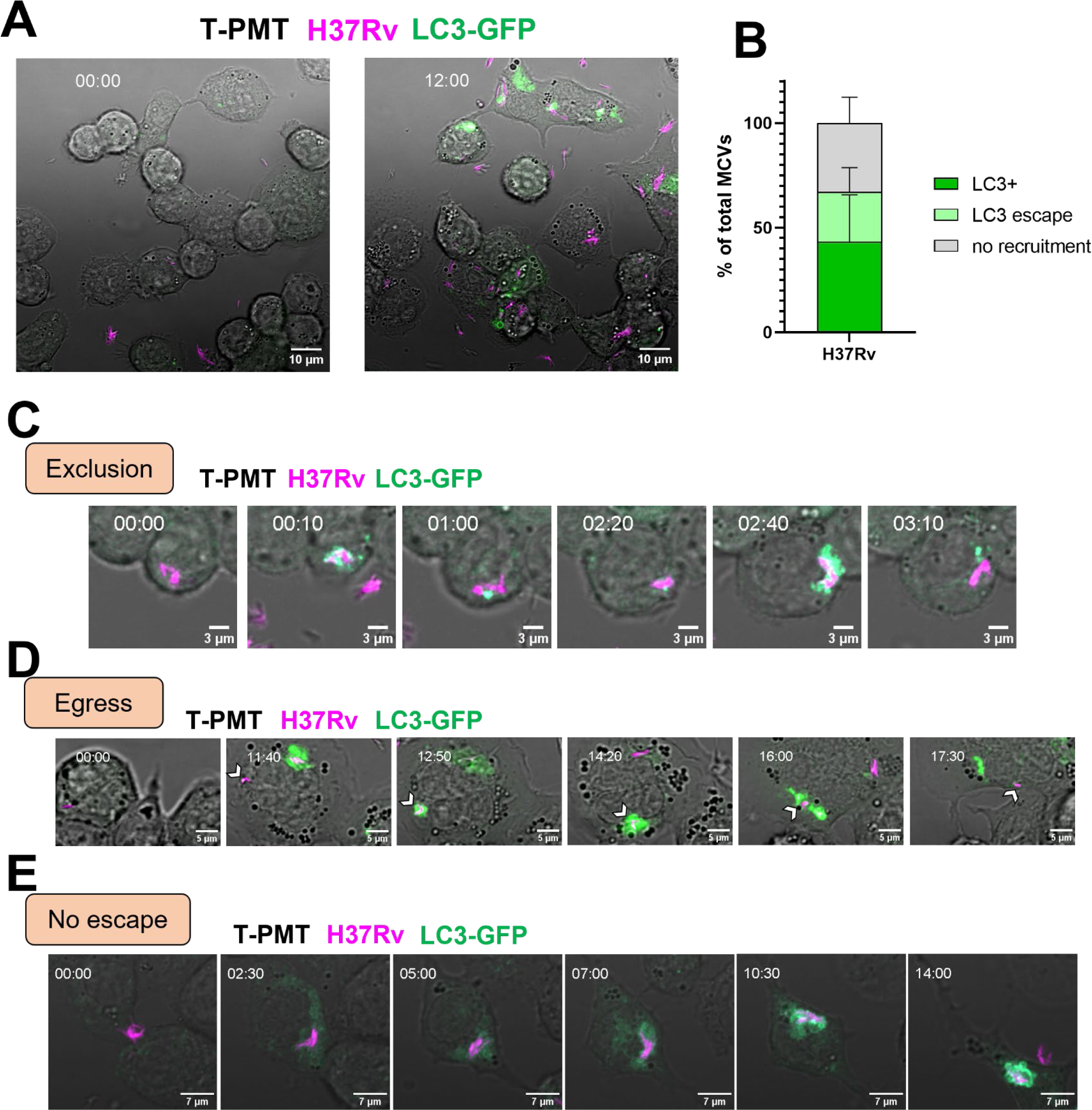
The Mtb phagosome induces frequent LC3 recruitment but can escape from it. THP-1 cells stably expressing LC3-GFP were infected with DsRed-expressing MtbH37Rv at an MOI of 2 for 16 – 20h. A 9μm z-stack with a 1μm step was recorded every 10 minutes. (A) A representative image of infected THP-1 cells at the beginning of recording and at 12h post-infection. Time stamp format is hh:mm. (B) quantification of LC3 recruitment to mycobacteria-containing vacuoles (MCVs) (LC3^+^), and subsequent LC3 escape or no LC3 recruitment. The graph shows the mean and standard deviation from 4 independent experiments, following between 22-43 MCVs per experiment. (C) Representative time lapse images extracted from Movie S2 of LC3 recruitment to an MCV followed by loss of LC3 signal by exclusion. (D) Representative time lapse images extracted from Movie S3 of LC3 recruitment to an MCV followed signal by egress of the bacteria out of the autophagosome. (E) Representative time lapse of an MCV exhibiting a LC3 recruitment

### No correlation between autophagy induction and the time after infection nor the size of the MCV

Given the dynamic nature of LC3-MCV interactions (Figure 1), we deepened the analysis of LC3 recruitment to MCVs. We analyzed a total of 73 MCVs showing at least one temporary LC3 recruitment by their time post-phagocytosis and their observed length of time of LC3 recruitment (table 1). Three groups of MCVs could eventually be distinguished: (i) MCVs showing LC3 recruitment between 0 and 1 h post phagocytosis (hpp) (Table 1, Figure 2B, C), (ii) MCVs showing recruitment after 1 hpp and (iii) the MCVs presenting multiple LC3 recruitment events. For each group, an example is given (Figure 2B–G), where the fluorescence quantification on the bacterial area over time (Figure 2C,E,F, Movies S4-6) could confirm the visual observation as shown on representative snapshots from the time-lapse study (Figure 2B,D,F). We observed a 40-minutes gap between early and late recruitment events, meaning that in the early event category LC3 recruitment was always observed between 0 to 0.33 hpp and the recruitment in the late event category started after 1 hpp (Table 1). However, within the late events group no clear pattern emerged that would indicate that the LC3 recruitment is governed by a clear timing factor. Instead, we observed a wide range of LC3 recruitment times, from 1 to about 13 hpp. indicating an absence of a coordinate response involving LC3 after Mtb phagocytosis.

**Figure 2:**
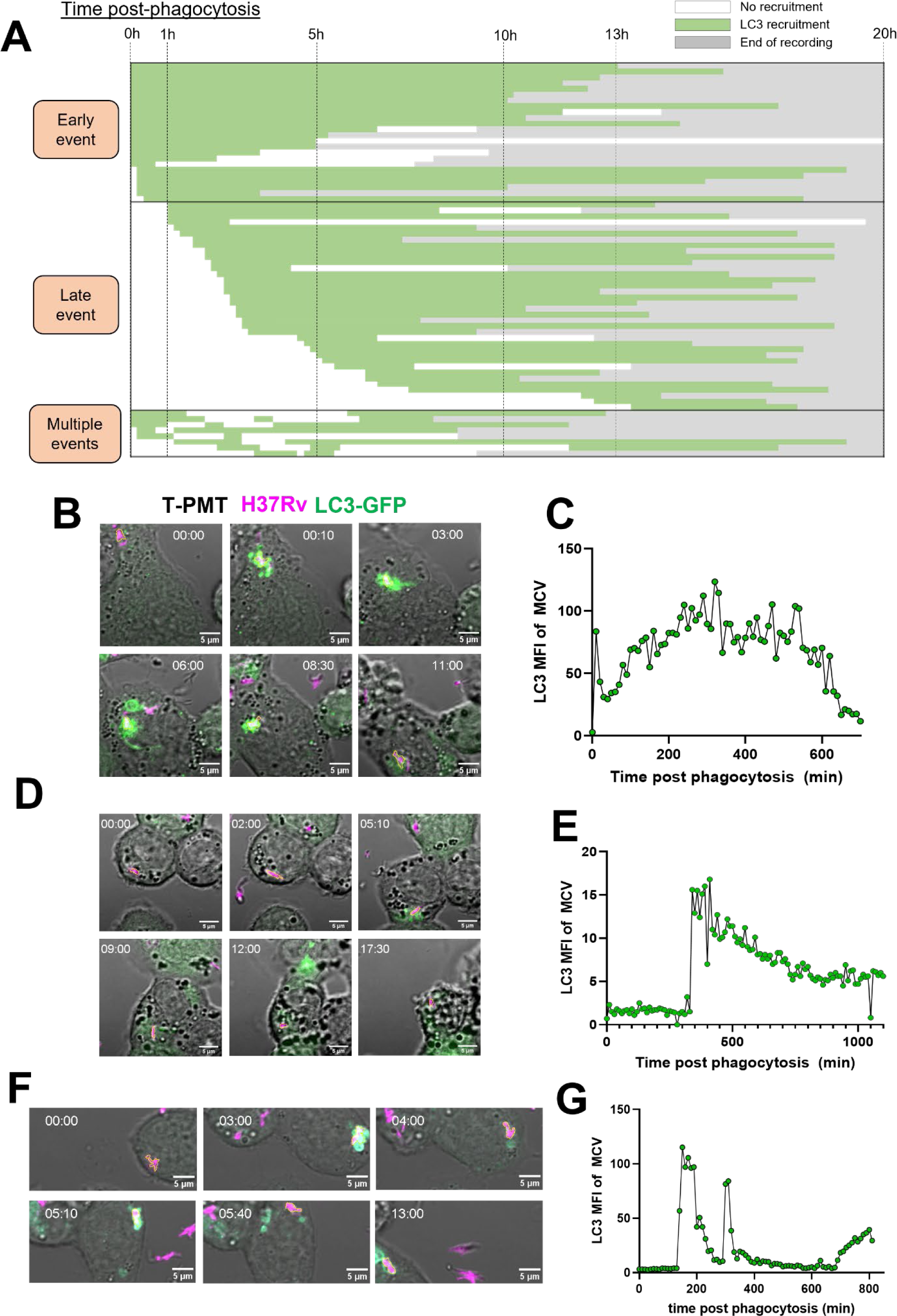
LC3 recruitment does not follow a specific timing pattern. (A) MCVs were categorized by their time of observed LC3 recruitment after phagocytosis. The green color indicates visually observed LC3 recruitment, and the white color no recruitment. The grey color indicates end of recording. The total number of MCVs observed was n=73. MCVs were organized into 3 categories: 1. “Early events” the ones showing LC3 recruitment immediately or shortly after phagocytosis (≤1h post phagocytosis), 2. “Late events” the MCVs showing an LC3 recruitment > 1h post phagocytosis and 3. “multiple events” the MCVs showing multiple LC3 recruitments. (B,D,F) Representative time lapses of MCVs of the 3 listed categories, extracted from Movie S4, S5 and S6 respectively. MCVs followed were outlines with a yellow region of interest (ROI) line. (C,E,G) Confirmation of visual recruitment by quantification of LC3 fluorescence intensity over time on the three MCVs in (B,D,F).

In summary, LC3 recruitment to MCVs is highly variable in timing, from early recruitments (< 0.33 hpp) to late recruitments (> 1 hpp), without clear patterns of a coordinated response.

A previous study proposed that the number of bacteria per MCV and more specifically their capacity to form cords could inhibit xenophagy in lymphatic endothelial cells.^51^ Intracellular cord formation is not occurring in differentiated THP-1 cells and thus could not be a factor in our system but the influence of bacterial burden per cell on LC3-recruitment was assessed. First, we compared the number of bacteria per MCVs to the time post infection when the first LC3 recruitment was observed (Figure 3A). A slight trend might be observed where the more bacteria per MCVs, the faster the LC3 recruitment seemed to be, but it failed to show any statistical correlation (R^2^= 0.072). This was confirmed by analyzing the average number of bacteria per MCV among the groups described in Table 1. On average, the number of bacteria per MCV was not different between the early and late recruitment groups or the group with multiple recruitment events (Figure 3B).

**Figure 3:**
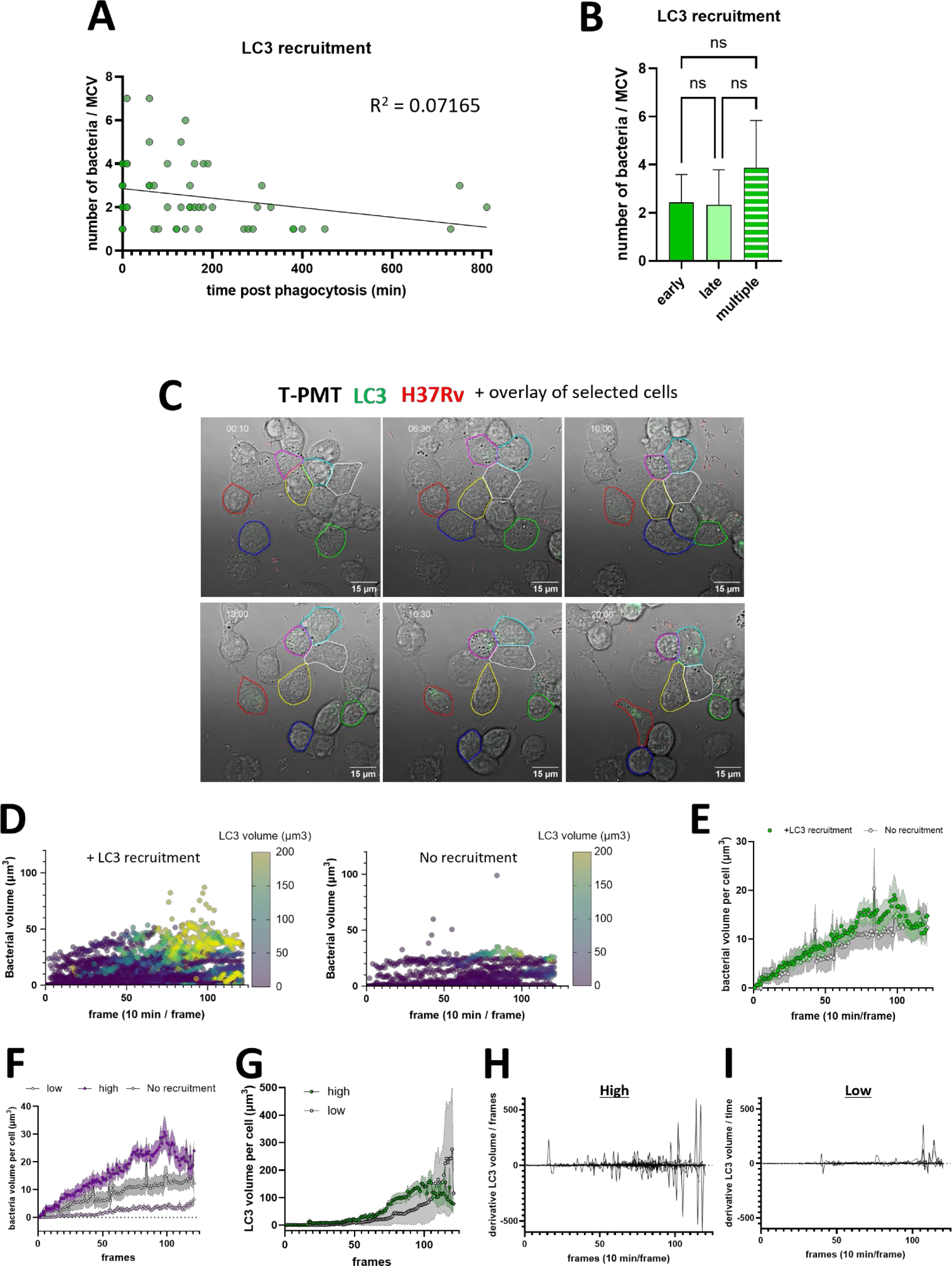
Macrophages bacterial burden does not directly influence the LC3 recruitment to MCVs. (A) Distribution of number of bacteria per MCVs compared to the time of observed of LC3 recruitment. (B) Quantification of the average number of bacteria per MCV in each group from figure 2A. (n=29 early event, n=36 late event and n=8 multiple events). The groups were compared by one-way ANOVA test followed by Dunnett *post-hoc* statistical test. (C) Representative time lapse with the overlay showing an example result of the single cell tracking, selection of cell of interest outlined and quantification described in methodology. (D) Single cell tracking and quantification of bacterial volume over time plotted on a 3 variables graph with the LC3 volume as a color dimension. Each dot represents the measurement in 1 cell in 1 frame. The left panel represents the quantification in cells exhibiting at least one temporary LC3 recruitment on at least one MCV (N_cells_ = 41, N_points_ = 4299). The right panel represents the quantification in cells that don’t exhibit LC3 recruitment (N_cells_ = 11, N_points_ = 1222). (E) Quantification of the average bacterial burden in μm^3^ over time, on the cell population described in (D). The dots represent the mean and the band around the connecting line the standard error to the mean. (F) Quantification of the average bacterial burden in µm^3^ over time, on the cell population exhibiting a high (>20 µm^3^, N_cells_ = 24) or low (<20 µm^3^, N_cells_ = 17) bacterial burden among the cells plotted in (D-left). (G) Calculation of the associated LC3 volume in the high and low cells. (H-I) The first derivative of the LC3 volume curves of each individual cells of the ‘high’ (H) or ‘low’ (I) groups was calculated and plotted. All the data here from 4 independent experiments.

### The bacterial burden does not directly drive LC3 recruitment but may accelerate its occurrence

Even though the size of the MCV itself did not appear to correlate with initiation of LC3 recruitment, in some infected cells, it seemed that subsequent LC3 recruitment happened because other MCVs were already present inside the cell. For example, in this recording of the cells (Figure 1D, Movie S3), the first bacteria entered the cell, and no LC3 recruitment to the MCV was observed for 11 hours and 50 minutes. Then a second bacterium was phagocytosed and immediately (≤ 10 min) triggered LC3 recruitment to that MCV. Interestingly, in this cell even the first MCV underwent LC3 recruitment shortly after the second MCV became LC3^+^. This series of events suggested that the second MCV triggered a response that then targeted both MCVs.

To analyze this phenomenon on a larger number of single cells, we implemented the new analysis method that we recently released^48^ on the different time-lapsed acquired. With this method, the cells that would stay in the field of view were tracked, isolated and separated into 3 groups: Uninfected, infected with LC3 recruitment on at least one MCV, and infected without visible LC3 recruitment on any MCV. The bacterial burden was calculated following the BBQ method recently released based on the relative quantification by bacterial volume calculation. To compare the level of bacterial burden over time with the level of autophagy induction, an analysis based on the volume of the LC3^+^ tubulovesicular structure was done based on the bacterial volume calculation logic (see methods). To validate that this approach is a relevant readout, the LC3 volume over time from single cells was pooled and plotted for the different cell populations (Figure S1). This confirmed that in the infected cells that visually showed LC3 recruitment, the LC3 volume consistently increased over time (Figure S1A), whereas in the infected cells without visual autophagy induction, no or barely and LC3 volume increase was measured (Figure S1B). Some increased LC3 volume increase was also occasionally observed in some uninfected cells (Figure S1C) but was associated with an increase in the overall signal of LC3 (Movie S1, uninfected cell in the center). From this, the bacterial volume was plotted over time with the measure LC3 volume as a 3^rd^ color dimension (Figure 3D). The increase in bacterial burden can be seen but seemed at least partially higher for the infected exhibiting LC3 recruitment. The displayed color for the LC3 volume seemed also in the brighter colors for the cells exhibiting a higher bacterial burden, meaning a higher LC3 signal in these cells. To deepen the analysis on the potential role of the bacterial burden in the induction of autophagy, the average bacterial burden over time between the 2 cells groups was calculated (Figure 3E). On average, there was no difference of burden between the cells with LC3 recruitment compared to the cells without. The absence of evidence from the influence of the bacteria may suggest that the deciding factor of autophagy may come from the host side, where a cell to cell variability may occur, like in the case of iNOS.^46^ We then decided to focus on the cells showing an induction of autophagy and hypothesized that in this population the burden may influence the autophagy dynamics. The cells were separated in two groups, the one reaching a bacterial volume ≥20µm^3^ (high) or <20 µm^3^ (low) (Figure 3F) and their respective average LC3 volume over time was plotted (Figure 3G). The results showed a noticeable trend on the LC3 recruitment being induced earlier in the high burden population compared to the low burden, before reaching an equivalent level. To visualize this phenomenon in a single cell fashion, the individual curves were transformed into their partial derivative calculated between each time points, plotted and superimposed by their respective cell category (Figure 3H,I). The derivative over time indicates the changes in the slop of the curve, thus should be able to display the sudden changes of LC3 volume that could happen in case of autophagy induction. Accordingly, in the cell population of high burden, the derivative showed a higher amplitude over time (Figure 3H), with a trend of LC3 volume changes earlier than for the cell population with low burden (Figure 3I). These results then strongly suggest that even if the burden is not a direct driver of LC3 recruitment, a higher burden may accelerate its occurrence.

### LC3^+^ MCVs do not acidify, and the recruitment frequency is not affected by IFN-γ pre-treatment of macrophages

During the time-lapses, a non-negligible number of bacteria did not show signs of escaping LC3 association (Figure 1B) suggesting that they might be contained in autophagosomes that could mature and eventually kill the bacteria. To explore the fate of LC3^+^ MCVs, the cells were infected for 4h and later dyed with Lysotracker blue (LTB) to assess phagosome or autophagosome acidity (Figure 4A). Strikingly, at 24 hours post-infection, none of the LC3^+^ MCVs were positive for LTB, although about 15% of MCVs were positive for LTB (Figure 4B). These results strongly suggested that LC3 recruitment does not result in Mtb contained within autophagolysosomes. Previous studies showed that pretreatment with IFN-γ increased LC3 association with MCVs and phagosome maturation.^4, 15^ To study this in a live-imaging condition, we opted for the far-red probe Lysoview-633 (LV), which showed a similar pattern to LTB at 24 hours post-infection and was not cytotoxic when left in the medium for 48 hours (Figure S2). The result of the pretreatment of THP-1-LC3-GFP cells with 10 ng/mL of IFN-γ overnight showed that the pretreatment did not affect the LC3 association level to MCVs and did not increase the phagosome maturation level at 6 hours post-infection, nor did it increase the number of acidified LC3^+^ MCVs (Figures 4D). Instead, the visual assessment of the recruitment indicated that Lysoview staining even tends to decrease (Figure 4D). To confirm these observations, we implemented a new workflow of high throughput single MCV fluorescence quantification (see methods, Figure S3). In the same trend, on average LC3 fluorescence on MCV was not different in cells treated or not with IFN-γ (Figure 4E, left). And the LV decrease was also confirmed in IFN-γ treated cells compared to untreated (Figure 4E, right). However, pretreatment with IFN-γ made the cells more susceptible to Mtb-induced cell death, characterized by “ballooning” morphology as observed through the transmitted light channel (Figure 4C, movie S9, S10). The quantification of this phenomenon at 20 hours post-infection showed an increase from 40% in untreated cells to 75% in IFN-γ pretreated cells after Mtb infection (Figure 4F). To explore the eventuality that this death could be due to difference in bacterial load or infectivity, these parameters were measured at 6h p.i. The results showed that no notable difference was observed between treated and untreated cells (Figure 4F). In summary, our results showed that LC3 recruitment almost never turned into a mature autophagolysosome, and that IFN-γ was not able to increase the frequency of LC3 recruitment or the maturation level of the autophagosomes.

**Figure 4:**
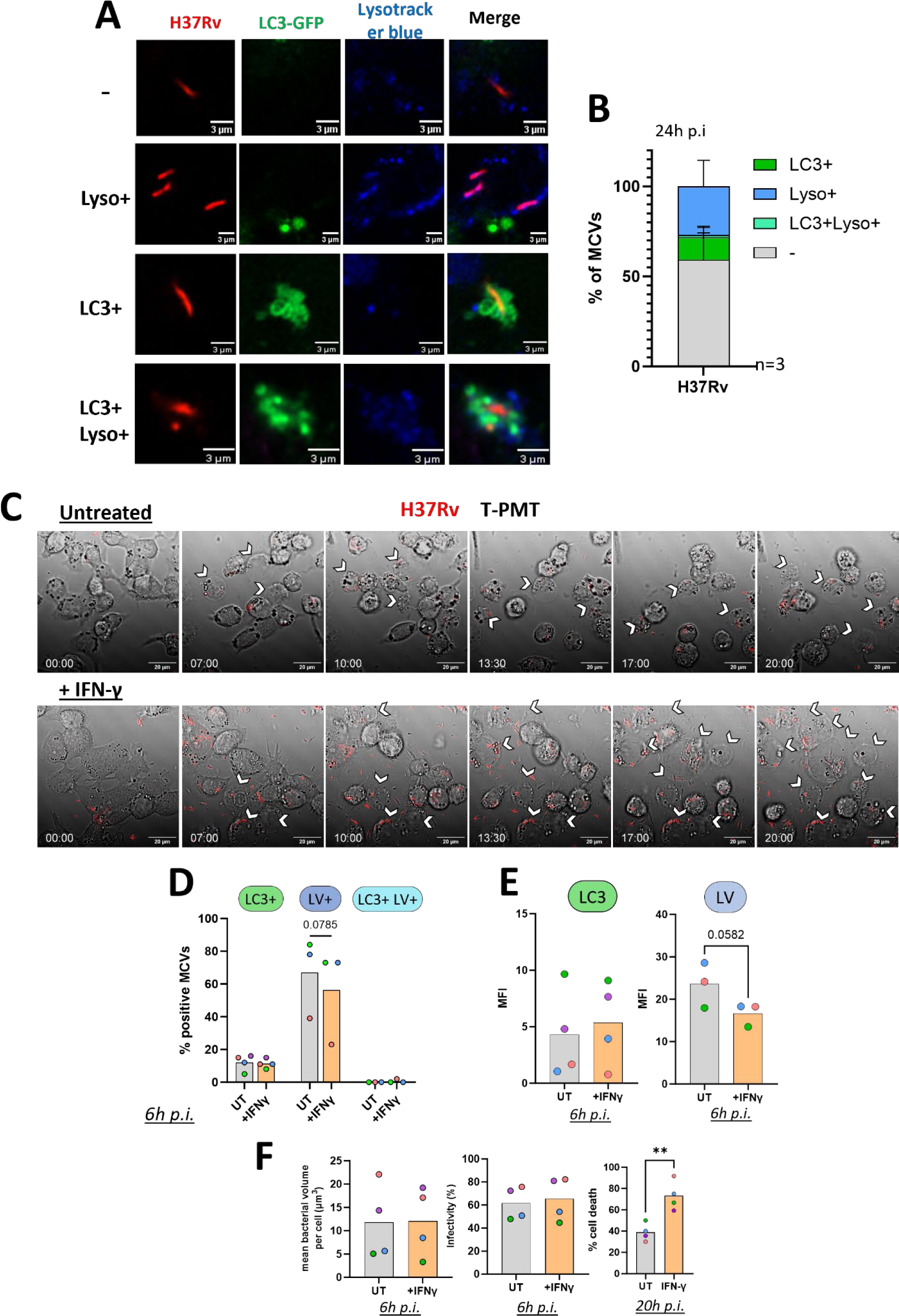
Autophagosomes are not bactericidal and IFN-γ treatment has no effect on LC3 recruitment frequency or phagosome maturation. (A-B) THP-1 expressing LC3-GFP were infected at an MOI of 10 for 4h and incubated for 24h. Cells were then stained with Lysotracker blue and imaged by confocal microscopy. (A) representative image of MCVs that were unstained, positive for LC3 alone (LC3+), positive for lysotracker alone (Lyso+), or double positive (LC3+Lyso+). (B) quantification of LC3 and Lysotracker staining on MCVs. The data were obtained from 3 independent experiments. (C-E) THP-1-LC3-GFP cells were treated with IFN-γ 10ng/mL or left untreated overnight and infected with DsRed expressing H37Rv at MOI 2. Cells were also stained with Lysoview-633 (LV) and the dye was kept during image acquisition. Cells were image by time lapse confocal microscopy. (C) Representative time lapse showing the untreated cells (top panel) or pre-stimulated with IFN-γ (bottom panel). Time stamp format is hh:mm. Chevrons are showing cells presenting the ballooning phenotype. (D) Quantification of the frequency of LC3 and/or Lysoview association to MCVs. (E) Quantification of the mean fluorescence intensity on single MCV in living cells. Data are from 3 to 4 independent experiments. The color of the points indicates data obtained from the same experiment. (F) Quantification of bacterial burden, infectivity using the volume calculation as described elsewhere. The cell death frequency was quantified by counting the proportion of cells exhibiting the ballooning phenotype during infection. Data are from 4 independent experiments. Points’ color is representative of the same experiment. Data between untreated and IFN-γ treated groups were compared by paired *t*-test. \*\**p*< 0.01.

### LC3 recruitment to and acidification of the MCVs are mutually exclusive events

Our steady state analysis showed minimal to none of the MCVs that were positive for LC3 and lysotracker staining (Figure 4B). From the untreated cells in figure 4C-E, we selected MCVs that showed LC3 recruitment and could be followed for at least 3h to investigate the dynamics of LV and LC3 association with the MCV. The analysis showed that LC3 and LV staining does not happen simultaneously (Figures 5A–D, Figure S4). Instead, we observed acidification of the MCV, followed by a drop of acidification prior to the subsequent LC3 recruitment. In some cases, the LC3 signal dropped and was followed by a quick recovery in acidification thus completing a cycle that could be repeated (Figures 5A, B, Movie S11). In other cases, the LC3 recruitment would persist for prolonged periods of time (>6h) until the end of the recording (Figures 5C, D, Movie S12). We did not observe cases in which the MCVs became LC3^+^ without first acidifying (LV^+)^. We also did not observe cases in which the LV^+^ MCVs lost acidification without gaining LC3 staining. To further visualize the link between acidification and LC3 recruitment, we plotted the LC3 signal from single MCVs against the LV fluorescence at every time point recorded (Figure 5E). The resulting dot plot and density map clearly show two populations of stained MCVs, the LC3^+^ ones, and the LV^+^ ones with very rare amount of double positive MCVs. This result confirms the observation that LC3^+^ phagosomes are rarely acidic and that an acidified vacuole loses its acidification before LC3 recruitment, indicating that membrane damages occurred and were recognized by the autophagic machinery. The cases of LV recovery concomitant with a LC3 loss of signal was also suggesting membrane repair involving the autophagy machinery.

**Figure 5:**
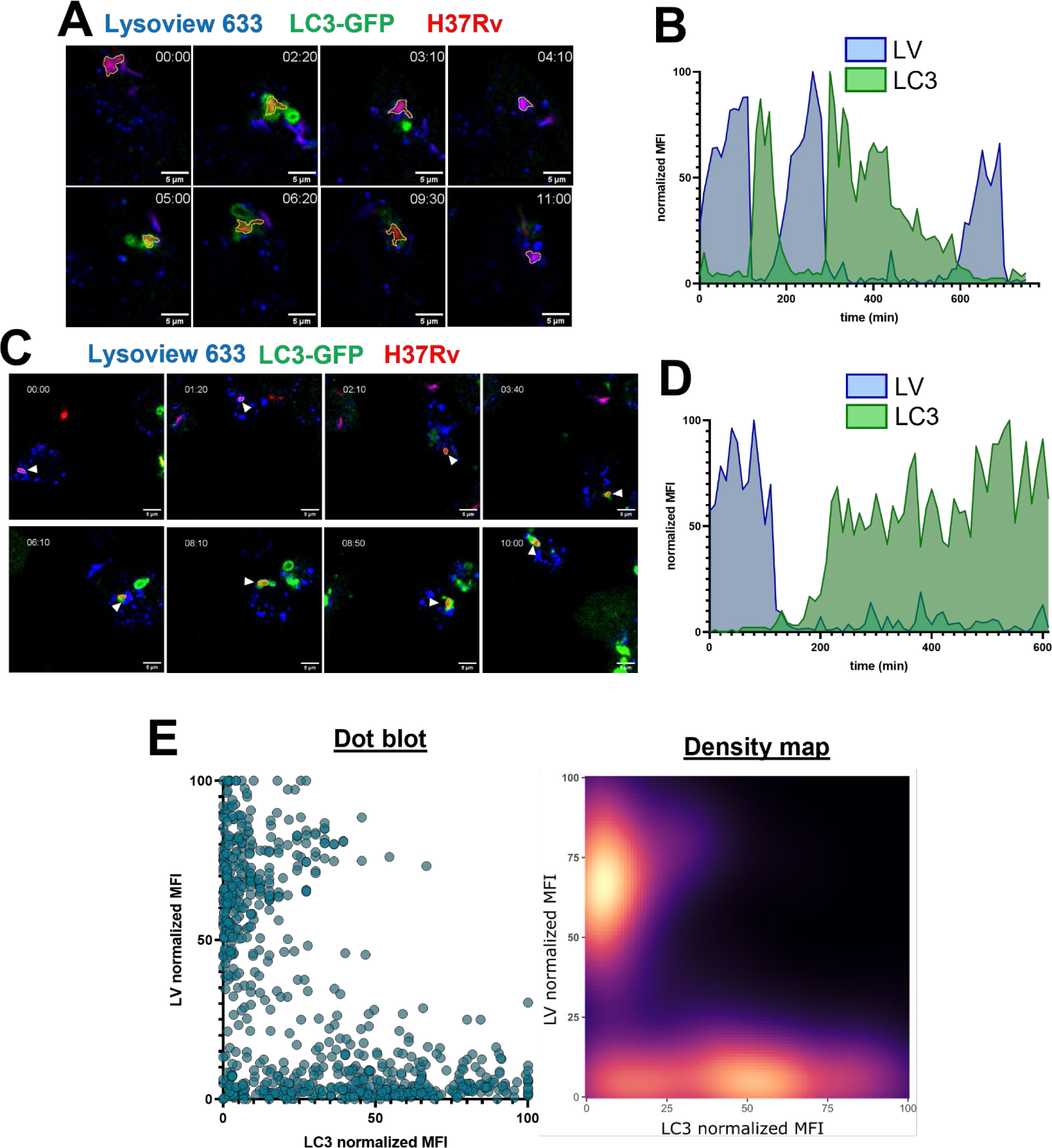
LC3 recruitment and phagosome maturation are mutually exclusive. Single MCV tracking in unstimulated THP-1-LC3-GFP cells infected with DsRed expressing H37Rv in figure 4. (A, C) representative time lapses of MCVs outlined with a yellow ROI. (B, D) quantification of fluorescence at the MCV for each time point. Mean fluorescence intensity of GFP and Lysoview was measured and normalized between values of 0 and 100 as 0 the minimum value recorded, and 100 the maximum value. (E, left panel) Quantification of Lysoview and LC3 normalized MFI and was compared. Each dot represents the GFP and LV normalized MFI at one time point for one MCV from 3 independent experiments (n=12 MCVs). The individual fluorescence traces of each MCVs are shown in Figure S4. (E, right panel) The distribution of data points in left panel is shown as a density map.

### Autophagy is associated with membrane damage and repair effectors but is not involved in membrane repair

Following the previous results of the dynamics of LC3 that points towards the detection of membrane repair and a potential involvement in membrane repair, we next evaluated the recruitment of known marker of membrane damage and repair that were shown associated with the MCV. Confocal immunofluorescence microscopy was used to detect Galectin-3 (GAL3), a known host cell marker of membrane damage induced by Mtb,^52^ ALIX, a core component of the ESCRT-III machinery that was recently found as a ligand of GAL3,^53^ and finally the oxysterol-binding protein 1 (OSBP) involved in ER-dependent membrane repair recently described in *Mycobacterium marinum* phagosome repair.^54^ At 20h post infection, the cells were stained for either of the markers and imaged by confocal microscopy. The results first confirmed that LC3 recruitment is in majority associated with GAL3 recruitment as previously described by others (Figure 6A-B).^45^ However, the LC3^+^ structures were rarely associated with the effector ALIX, showing that ESCRT-III machinery was not recruited in these experiments (Figure 6A-B). This result prompted the investigation into ER-dependent membrane repair that was found involved in mycobacterial infections^54^. Accordingly, LC3^+^ MCVs were in majority co-stained with OSBP (Figure 6A-B), thus showing that LC3 recruitment is indeed associated with membrane repair and that the acidification recovery observed previously (Figure 5A-B, S4) corresponds to the phagosomal membrane getting repaired, likely through OSBP activity.

**Figure 6:**
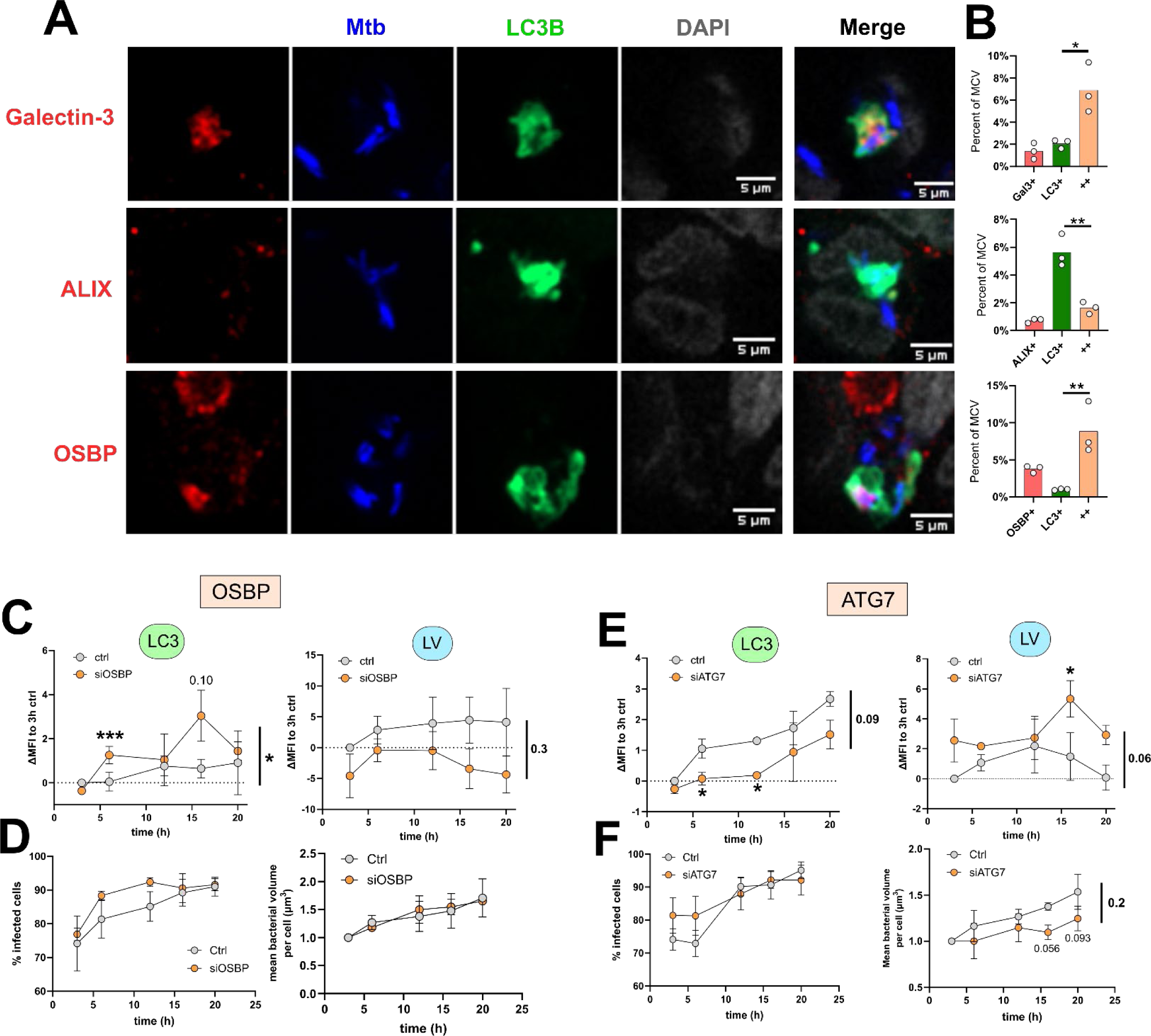
Autophagy on the MCV correlates with phagosome damage and ER-dependent membrane repair, but does not contribute to membrane repair. (A-B) THP-1-LC3-GFP cells were infected with DsRed expressing H37Rv (Mtb) at an MOI of 10 for 4h, chased and fixed after an incubation of 20h. The cells were stained either for Galectin-3, ALIX or OSBP and imaged by confocal microscopy. (A) Representative images of MCVs. (B) Quantification of the frequency of association of the markers to the MCV. Points represent the averages of the quantification on 465-1558 MCVs obtained from 3 independent experiments. The difference between LC3+ and ++ conditions was individually analyzed by paired t-test. \**p* ≤ 0.05, \*\**p* < 0.01. (C-F) THP-1-LC3-GFP cells were transfected with siRNA against OSBP (C-D) or ATG7 (E-F), infected with Mtb, stained by Lysoview-633 (LV) and imaged by time-lapse confocal microscopy. The movies were analyzed to measure the mean fluorescence intensity (MFI) of LC3-GFP and LV on single MCV in living cells at the designated time points (C, E). The cells infection rate and the intracellular growth were calculated at the designated time points (D,F). The curves represent the difference of the mean fluorescence intensity (MFI) to their correspondent uninfected 3h time point, from 3 independent experiments. Each mean was obtained from the measurement on 99-358 MCVs. The difference between time points was individually analyzed by paired t-test, and the curves as a whole were analyzed by modified chi-squared method (see methods).

The next question was to determine if autophagy and LC3 recruitment were directly involved in the membrane repair that was previously observed. To answer this question, ATG7 and OSBP (as a positive control for membrane repair) were respectively knock-downed using siRNA (Figure S6), and the cells infected with Mtb and followed by time-lapse confocal imaging. The expected result was that a direct involvement of autophagy in membrane repair would decrease the average signal of lysoview on the bacteria as there would be accumulation of damaged and unrepaired phagosomes. As for OSBP knock-down (KD), the expected observation would also be accompanied with an increased LC3 recruitment as an accumulation of damaged compartment would increase the frequency of autophagy induction around bacteria. Thus, the mean fluorescence intensity of LC3 and lysoview was measured on single bacteria at different time post-infection (Figure C, E). For OSBP-KD, as expected the silencing resulted in at least a temporary increase in LC3 recruitment on the MCVs, while the LV tended to decrease (Figure 6C). However, in the ATG7-KD condition, while the silencing reduced LC3 recruitment as expected, the lysoview staining was at least temporarily increased on the MCVs (Figure 6E). To alleviate any effect due to bacterial burden (figure 3D), the effect of the respective KD on bacterial burden and infectivity was measured (Figure 6D,F, Figure S7). The result showed no clear differences in intracellular bacterial growth nor the infectivity over time in the OSBP-KD cells and the ATG7-KD cells compared to control, or even a decrease in growth in the ATG7-KD (Figure 6D, F). This demonstrates that autophagy is not directly involved in the membrane repair observed. The increase in LV MFI (Figure 6E, right) could even indicate that autophagy would compete with the different membrane repair machineries at play.

## Discussion

Our study is the first in depth quantitative analysis of membrane damage and autophagosome formation on single MCVs over time during Mtb infection. Our results and published data are summarized in the model shown in Figure 7 which focuses on the MCV and the observed role of LC3 recruitment. We show that autophagy recruitment to the MCV is a very dynamic event without any clear timing pattern post-phagocytosis and no correlation with the bacterial burden. As published before the induction of autophagy is linked to membrane damage, but our novel observation is that its activation on the MCV is reversible as the signal could disappear likely in correlation with a membrane repair phenomenon but could eventually be triggered again in a cycle of damage and repair of the phagosome.

**Figure 7:**
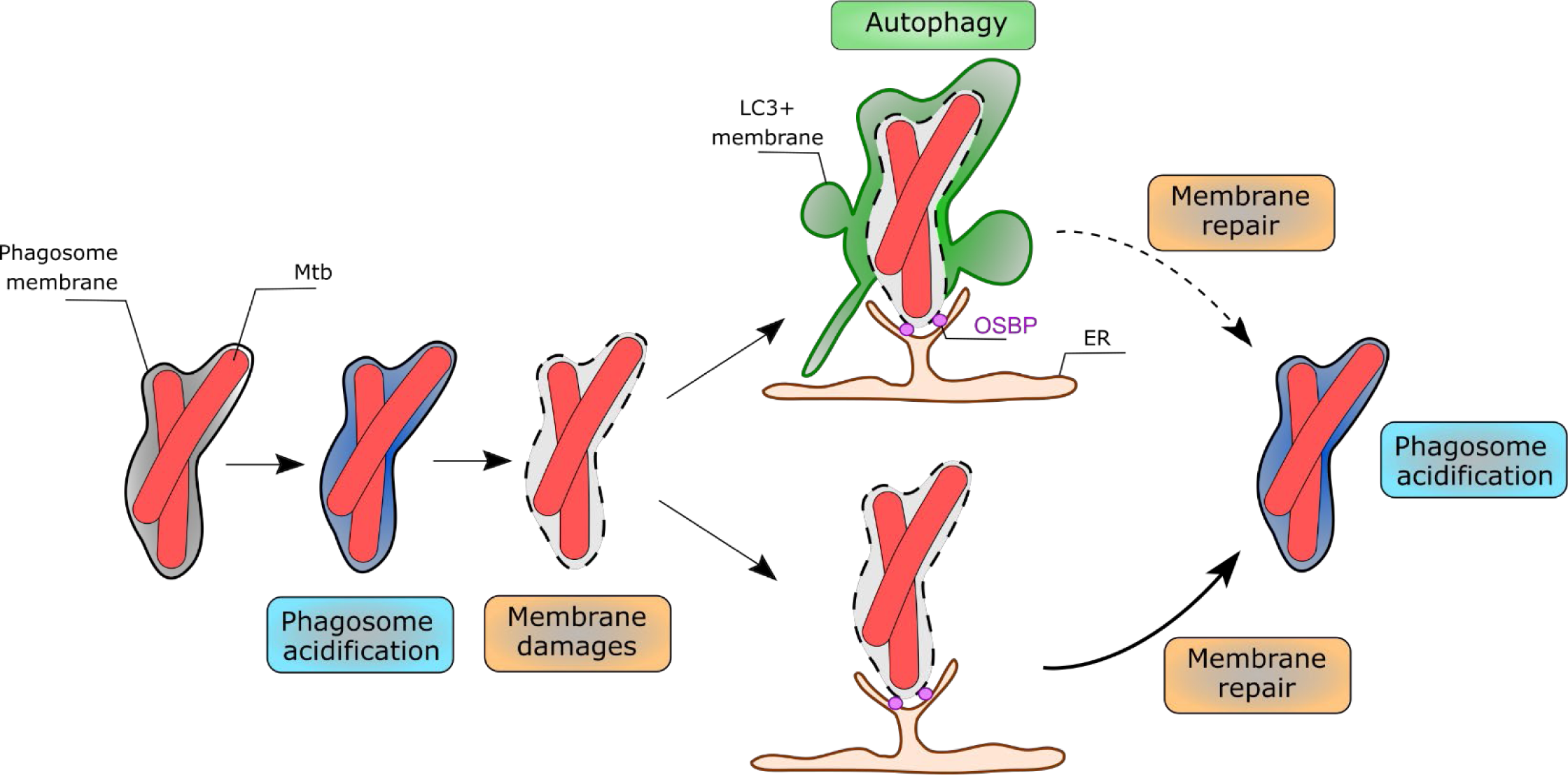
Model of LC3 recruitment to the Mtb phagosome. After phagocytosis, the MCV may undergo acidification. This acidification is lost upon induction of membrane damage which triggers LC3 (autophagy) and OSBP recruitment. An ER-but not autophagy-related membrane repair response eventually allows the rescue of acidification of the MCV but can subsequently be damaged again. Some bacteria are eventually able to physically escape from the phagosome or the LC3 positive vacuole to reside in the cytosol.

The loss of acidity of the MCV as reported by the loss of lysotracker/lysoview staining is also an indicator of membrane damage as it was used to follow lysosome damage and repair mechanisms via live cell imaging.^55–58^ Importantly, this method has been used to assess phagosomal membrane damage during Mtb infections as well.^33, 59^ The time-lapse analysis shows that acidic MCVs lose lysoview staining right before they recruit LC3 (Figure 5,7). The rapid and subsequent LC3 recruitment / autophagy observed here has also been previously associated with membrane damage.^16, 17, 33, 45^ To definitively confirm the presence of membrane damages, the cells were stained for galectin-3 and the immunofluorescence results demonstrated that LC3 recruitment is accompanied in majority by galectin-3 recruitment (Figure 6A-B). Previous work on the dynamics of galectin-3 and LC3 recruitment suggested that galectin-3 recruitment preceded LC3 recruitment,^45^ and accordingly, galectin-3 can stimulate the activation of autophagy.^53^ The damages can be explained by the EsxA/ESX-1-induced phagosomal membrane damage described by many other studies, leading to loss of acidity. The recovery of the acidification after damage then represents membrane repair as the restoration of acidity requires an intact membrane. In that sense, the autophagosomes were also in majority co-stained with the ER-dependent membrane repair effector OSBP, but not with ESCRT-III machinery effector ALIX (Figure 6A-B). This result confirms a recent study demonstrating the activation of ER-dependent membrane repair in the *Dictyostellium discoidium*-*Mycobacterium marinum* model system.^54^ The OSBP recruitment was confirmed in human macrophages during Mtb infection.^54^ The lack of observation of ALIX recruitment suggested that the ESCRT-mediated repair machinery is not involved in the repair is unexpected. An explanation may be that the implication of galectin-3 to recruit ALIX and initiate ESCRT-mediated repair was observed in different cell types, mainly in HeLa cells.^53^ It is conceivable that in THP-1 cells the primary repair mechanism involved is the ER-membrane repair mechanism rather than ESCRT-III due to differential expression of the factors. Moreover, these studies mainly use Leu-Leu methyl ester hydrobromide (LLOMe), whose mechanism of action to disrupt membranes may differ from the damages induced by Mtb, which mechanism is not entirely resolved. Additional experiments would be required to definitively rule out the involvement ESCRT-related effectors, in the repair of damaged MCVs in our system.

As our data shows that autophagy signal disappeared with the membrane being repaired demonstrating an unexpected reversible property of autophagy and autophagosome formation. This also begged the question of a potential involvement of autophagy in repairing the membrane. This hypothesis could be in agreement with the role of LC3 in phagosome membrane repair during Salmonella infections^60^ and a similar proposed mechanism during *Mycobacterium marinum* infections.^24, 25^ ATG7 and ATG14 KO in human macrophages resulted in a higher fraction of Mtb bacteria in the cytosol, suggesting that autophagy could contribute to membrane repair and keep the bacteria in a phagosome.^12^ However, in our experiments, ATG7 KD exhibited opposite effects to OSBP KD, demonstrating that in our cell system autophagy is not directly involved in the repair (Figure 6). The increase in lysoview staining even suggests that autophagy may actually delay the repair mechanisms maybe by physically competing with access to the MCV (Figure 6,7). An explanation could be that both autophagy and membrane repair machineries both are activated at the site of membrane damage but that their function is different. Indeed, autophagy is crucial for the removal and recycling of intracellular components, whereas ER-dependent membrane repair machinery restores the compartments integrity.^58^ LC3 recruitment to the MCV is dependent upon the Mtb ESX-1 secretion system and requires damage to the phagosome membrane since it was observed galectin-8 recruitment precedes LC3 recruitment to the MCV.^5, 16, 33^ We observed that MCV acidification frequently precedes LC3 recruitment (Figure 5). The MCV acidification was suggested as a trigger for Mtb to induce membrane damages. Indeed, the Mtb transcriptional regulator PhoP/R system is activated by slightly acidic pH^61^ and PhoP/R is a positive regulator of EsxA secretion.^62^ In addition, the membrane lytic activity of EsxA is increased by an acidic pH possibly via the pH-mediated dissociation of EsxA from its chaperon EsxB.^63–65^ Hence the acidification of the MCV we observe could lead to increased levels of active EsxA within the phagosomal lumen which results in phagosomal membrane damage leading to loss of protons and hence lysoview staining. However, it was also shown that phagosome maturation blockade is a prerequisite for Mtb phagosome escape^66^ and that blocking of phagosome acidification with bafilomycin strongly promotes phagosome membrane damages.^66, 67^ Accordingly, our data shows that many MCVs did not lose acidification and persisted in this environment for the whole recording (example Movie S13). Thus, the requirement of acidification to induce membrane damage to the MCV is still an open question. Additionally, the lysoview probe used in this study can emit fluorescence at pH ≤ 7 according to the manufacturer (Biotium; Cat#70058). It was observed that the phagosome maturation blockade by Mtb stabilize the pH at ∼ 6.5,^35, 68^ which could still potentially be stained by lysoview in our experimental conditions. Thus, we cannot firmly differentiate highly acidic compartments from the phagosome where Mtb at least partially blockaded the maturation and consequently additional studies would be required to conclude on the matter.

The final aspect of this LC3 recruitment study was to investigate how potentially bactericidal autophagy could be by looking at the acidification level of the phagosome/autophagosome in combination or not with an IFN-γ stimulation that is supposed to enhance the phagosome/autophagosome maturation.^69^ In contrast to other studies in BMDMs,^15^ IFN-γ pretreatment did not affect LC3 recruitment and phagosome acidity at 6 hours post-phagocytosis in THP-1 cells. This results shows a potentially crucial difference between mouse and human macrophages and confirms what was reported elsewhere that IFN-γ had no noticeable effect on Mtb fitness in primary human macrophages^4, 33^ and phagosome maturation.^4^ However, in our study, IFN-γ had a strong effect on macrophage viability during infection (Figure 4F). This is most like due to a pathway described in a study showing that stimulating human primary macrophages with IFN-γ promotes a necrotic cell death characterized by DNA release during Mtb infection.^70^ Alternatively, our result might be caused by a similar cell death mechanism observed in macrophages stimulated with LPS and IFN-γ, which was able to cause a necrotic type of cell death which is not associated with a protective host response to Mtb infections.^71^ In the experiments where ATG7 was knocked-down, the monitoring of bacterial burden over time did not show that the reduction of autophagy increase bacterial growth, reinforcing the idea that autophagy is not actively killing the bacteria (Figure 6). A limitation of these experiments is that the duration of the recording is only about 20h which may not be enough time to observe differences in growth rate. However, in our continuous infection model, the cell death level of THP-1 cells beyond 24h rapidly got extremely high which prevented a more thorough analysis at later time point post infection (data not shown). A previous study showed only a difference in growth of Mtb between WT and ATG7-KO in human macrophages at 72 or 96h p.i.^12^ Additionally, the growth curves provided in this study show a delay of growth by about 24h rather than a killing of the bacteria.^12^ This study and our results support a model in which autophagy only has a modest role as a bactericidal host defense mechanism in human macrophages. The impact of xenophagy seem to be more striking in primary murine macrophages with a severe delay in growth of Mtb in WT macrophages compared to *Atg5, Atg7* or *Atg16L1* KO cells).^72^ The deletion of genes of proteins involved in canonical autophagy (such as ATG5) leads to increased susceptibility of the mice to the infection.^72^ However, the mechanism underlying the susceptibility may not be linked to cell autonomous defense but possibly more related for example to a role of ATG5 in immune response modulation during the infection such as in the promotion of neutrophils recruitment.^13, 73^ A follow-up study by the same group also observed that autophagy does not limit Mtb replication in macrophages in *vivo*.^14^

The LC3 recruitment by the LC3 lipidation on single membrane is proposed to be named atg8ylation and could be mobilized during diverse membrane associated events.^74^ The only atg8ylation phenomenon described so far in the context of Mtb infection is LAP.^15^ Our study failed to observe signs of LAP being induced after Mtb phagocytosis as the events of LC3 recruitment showed tubulovesicular structure formation, which is induced during canonical autophagy or xenophagy and is distinctively different from LC3 lipidation at the phagosome membrane.^75^ We can propose a hypothesis for the lack of detection of LAP in our work. First, Mtb inhibits LAP through the NOX2-inhibiting activity of the Mtb CspA protein^15^ and this inhibition might be more potent in human macrophages when compared with BMDMs, in which most of the published work has been performed to date. Another explanation could be the dependency of ROS production for triggering LAP. Indeed, Mtb possesses multiple factors that can neutralize the ROS produced, like KatG^76^ and SodA,^77^ and other factors linked to a decrease in ROS production, such as NuoG.^78^ Another study also found that PPE2 can bind p67, a subunit of the NOX-2 complex, to prevent ROS production.^79^ In summary, all these factors could prevent LAP and the associated LC3 recruitment. Also, we did not detect MCVs that were acidified and LC3^+^ and a recent study found that LAP was dependent on V-ATPase recruitment to the phagosome and that the stabilization of the complex by saliphenylhalamide further stimulated the LAP and LC3 recruitment.^80^ Hence, we think that the LC3 recruitment to the MCV in our study was almost exclusively associated with an autophagy-related membrane damage repair mechanism and/or autophagosome formation.

Our experimental approach allowed us to determine at what time after phagocytosis MCVs become LC3^+^ and we noticed that there was a wide range of times post-infection that this event could occur (table 1). Thus, we investigated if LC3 recruitment could be linked to the number of bacteria per MCV by tracking the bacterial fluorescence signal per MCV and the LC3-GFP signal over time. Our results showed that there was no correlation between the number of bacteria per MCV and the time of LC3 recruitment (Figure 3A). Quantification of the average number of bacteria per MCV exhibiting early recruitment (< 1 hour) or late recruitment (> 1 hour) also demonstrated no difference (Figure 3B). To the best of our knowledge this is the first time that a detailed analysis and quantification of these parameters was done since the only other time-lapse study following LC3 recruitment onto MCVs aimed to study the structure of LC3^+^ MCV by CLEM and FIB-SEM imaging^33^. We observed that an MCV failed to show LC3 recruitment for 10 hours but became LC3^+^ soon after another bacterium was phagocytosed into the same cell. So, we hypothesized that the existence of an MCV prior to the phagocytosis of a new bacterium could trigger LC3 recruitment to all MCVs, or that the bacterial burden could directly influence LC3 recruitment. To test the hypothesis, we implemented the new workflow we recently released^48^ to analyze the correlation between the activation of LC3 recruitment and the bacterial burden at the single-cell level over time. By this approach, we observe no difference of bacterial burden over time between cells that show at least one LC3 recruitment to MCVs compared to the one that did not (Figure 3E). In the eventuality that the LC3 recruitment is also variable on a cell-to-cell basis, the analysis was refined only on the cells showing LC3 recruitment to see if in these cells, the bacterial burden may have influenced the LC3 recruitment (Figure F-I). Indeed, the cells having a burden higher than 20μm^3^ showed a clear trend for an earlier autophagy induction than the cells having a burden lower than 20μm^3^ (Figure G-I). This could be explained now in regard to the induction of membrane damage, where an increasing amount of bacteria may induce more damage, pushing the cells toward an induction of autophagy. It was also observed that Mtb infection can promote the expression of autophagy effectors,^81^ thus an increasing number of MCV may even promote this expression further.

Our study demonstrates a direct implementation of the workflow we recently released^48^ for high throughput, single cell analysis of infected cells dynamics, and single MCV quantification of marker recruitment / presence. By applying this workflow to our time-lapse datasets, we were able to track and select different cell populations with distinct observable behaviors, and measure parameters of interest in a automated and limited bias fashion. This was done using open-source and free software that demonstrates the feasibility without having to rely on costly licensed software. As a statement of its flexibility, we also adapted the single cell analysis workflow to automatize the measure of fluorescence at the MCV to provide a limited bias quantification of events than manual scoring. This was done due to the mobility and amount of MCV per cell preventing the direct use of the single MCV tracking and quantification workflow. This approach allowed to confirm the visual assessment of that the treatment of the cells with IFN-γ had to effect to a decrease in lysoview staining, but no difference in LC3 recruitment. This method was also used on the time-lapse experiments, sufficient to reveal the dynamics of phagosome maturation and autophagy in the context of ATG7 and OSBP KD. Based on our results, the expression of fluorescent LC3 combined with lysoview also provides direct observation of membrane damages induction and membrane repair in a time-lapse microscopy set-up. Thus, these types of markers combined with time-lapse microscopy could be a valuable tool for the future study of membrane repair mechanics and dynamics, during infection of macrophages or other host cells by intracellular pathogens.

## Material and methods

### Reagents and antibodies

Lysotracker blue (Molecular Probes), Lysoview-633 (biotium), Phorbol-Myristate-Acetate (PMA, provider), RPMI medium (Gibco), Fetal Bovine Serum (FBS, Gibco), DPBS 1X (Gibco), Recombinant human Interferon Gamma (R&D Systems), OADC supplement (R&D Systems), Zeocin (invivogen), anti-ATG7 antibody (Cell signaling technology E6P98), anti-OSBP antibody (Sigma-aldricht HPA039227), anti-Galectin-3 antibody (Cell signaling technology D4I2R XP(R)), anti ALIX antibody (Santa Cruz Biotechnology, sc-53540), Goat anti-mouse AF647 antibody (Jackson laboratory). Goat anti-Rabbit AF647 (Jackson Laboratory), Goat anti-Rabbit HRP (Jackson Laboratory), Goat anti-mouse HRP (Jackson Laboratory).

### Mycobacterial strain and culture

The DsRed-expressing strain of *M. tuberculosis* H37Rv was generated by cloning the *dsred* gene from the pMSP12-dsred2 (addgene #30171) into pMAN-1 expression vector previously used^82^ by EcoRI and HindIII digestion. The *dsred* gene was amplified using the following primers sequences:

Forward CGGCGAATTCATGGCCTCCTCCGAGAACG;

Reverse GGCTAAGCTTCTACAGGAACAGGTGGTGG

Bacteria were grown in liquid Middlebrook 7H9 medium supplemented with 10% oleic acid-albumin-dextrose catalase (OADC) growth supplement, 0.2% glycerol, 0.05% tween 80, and Zeocin 100µg/mL. For infections, the bacteria were washed in 0.05% PBS-tween 80 before being added to the infection medium (see procedure below).

### Monocyte culture, differentiation and infection

GFP-tagged LC3 expressing THP1 monocytes (THP-1-LC3-GFP) were provided by Dr. John Kehrl (NIH) and used in a previous study.^49^ The cells were cultured in RPMI 1640 (ATCC modification) supplemented with 10% heat-inactivated Fetal bovine serum (complete medium). The cells were differentiated with 20 ng/ml of PMA for 20–24 hours. In the case of IFN-γ pre-stimulation, the cells were washed in 1X DPBS and incubated overnight with 10 ng/mL of IFN-γ in complete medium. Stimulation efficiency was evaluated by flow cytometry and by the staining of MHC-II (HLA/DR) surface expression (Figure S3).

For infections, cells were washed in 1X DPBS and incubated in RPMI supplement with 5% normal human AB serum (infection medium). The bacteria were added to the cell at an MOI of 2 without a washing step for time-lapse imaging or an MOI 10 for 4 hours for single time point imaging. After the 4-hour incubation, cells were washed and incubated for 16 hours in complete medium supplemented with 100 μg/mL Gentamycin.

### Knock-down of ATG7 and OSBP

The procedure for the nucleofection of siRNA was done following the neon transfection manufacturer’s instructions.

The THP-1 cells were differentiated as described above and then detached using the Cell stripper (Corning) solution. The cells were then pelleted and resuspended in the E2 buffer that is part of the Neon transfection reagents kit (thermofisher). 100pmoles of siRNA for ATG7 (#ref) or OSBP(#ref) was added and the cells were electroporated using 2 pulses, 1700V. The cells were incubated in complete culture medium for 48 hours in glass-bottom observation chambers (ibidi) at 2×10^5^ cells/well.

### Western Blot

After the 48h of incubation, an amount of 5×10^5^ of transfected cells were lysed with lysis buffer (1% NP-40, 0.4 mM EDTA, 10 mM Tris-Hcl, 150 mM NaCl) for 2 minutes at room temperature. The proteins were dosed using the BCA protein dosage kit (thermofisher) and according to the manufacturer protocol. Then 10 µg of lysate was mixed with Laemmli buffer 1X (BioRad) with β-mercaptoethanol 10% and boiled at 95°C for 5-7 minutes. The proteins were separated by SDS-PAGE in a 12% polyacrylamide gel (Genscript) at 200V for 30 min. The proteins were then transferred on a nitrocellulose membrane. The membrane was then washed with DPBS + tween20 0.1% (PBST) and block with PBST-milk 5% for 30 min at room temperature. The membrane was then incubated with PBST-milk 0.5% + anti-OSBP (1:1000) or anti-ATG7 (1:1000) antibody + anti-β-actin (1:5000) at 4°C overnight. The membrane was washed with PSBT three times and then incubated with PBST-milk 0.5% + goat anti-mouse HRP 1:15000 + goat anti-rabbit HRP 1:15000 for 2h at room temperature. The membrane was washed three times with DPBS and revealed using with SuperSignal West Pico ECL kit (####, Thermo Scientific).

### Time-lapse confocal microscopy

THP-1 cells were seeded and differentiated in glass-bottom observation chambers at 2×10^5^ cells/well. The cells were stimulated with IFN-γ, if needed, before the infection (see above). For imaging of phagosome maturation, the infection medium was supplemented with Lysoview-633 at a final concentration of 1:2000 in infection medium and left on the cells during recording. The cells were imaged using a Zeiss laser scanning confocal microscope LSM-800, equipped with two gallium arsenide phosphide photomultiplier tube (GaAsP-PMT) detectors and a transmitted light photomultiplier tube detector (T-PMT), using the 63x/NA1.4 Oil objective. The observation chamber was maintained at 37°C and supplied with 5% CO2 through a humidifier system. A 10 μm z-stack (1 μm steps) was acquired every 10 minutes on 4 to 8 fields of view over 16 to 20 hours.

To assess phagosome maturation using lysotracker blue, the cells were infected in glass bottom observation chambers at an MOI of 10, as described above, and incubated for 16 hours. The cells were washed three times with 1X DPBS and incubated in complete medium supplemented with 1:10000 lysotracker blue for 30 minutes. The cells were washed, incubated in complete culture medium, and imaged by confocal microscopy.

### Immunofluorescence and confocal microscopy

The THP-1-LC3-GFP cells were seeded in a 24 wells plate with glass coverslips at the bottom, at a density of 5×10^5^ cells / well. The cells were differentiated as described and infected for 4h at an MOI of 10. The cells were washed three times with DPBS and incubated in complete medium supplemented with Gentamycin at 100 µg/mL for 20h. Then, the cells were washed three times with DPBS and fixed in PFA 4% for an hour at room temperature. After fixation, the cells were washed three times with DPBS and incubated twice for 5 minutes with DPBS supplemented with Glycine (100mM). The cells were washed once with DPBS and then blocked with DPBS + BSA 0.2% + Saponin 0.05% (PBSAP) + FcBlock TruStain 1:100 (BioLegends) for 30 minutes. The cells were incubated anti-Galectin-3 (1:200), anti-OSBP (1:80), or anti-ALIX (1:200) in PBSAP at 4°C overnight. They were then washed three times with PBSAP and then incubated with PBSAP + goat anti-mouse or goat anti-rabbit AF647 antibodies (1:200) for 2h at room temperature. The cells are finally washed with PBSAP three times and then mounted on slides using Prolong gold anti-fade with DAPI. The cells were imaged on a Zeiss LSM 980 Airyscan 2 confocal microscope equipped with a 63X oil-immersion objective.

### Image analysis

All the MCV qualitative assessments of LC3 recruitment, fluorescence quantifications, single cell tracking and fluorescence quantification, and single MCV fluorescence quantification were performed using the ImageJ/Fiji software^83^ directly using custom macros or in python using the library PyimageJ.^84^

The method for manual tracking and quantifying the fluorescence at the MCV over time in Figure 5 was previously described.^85^ For the single cell tracking and quantification of bacterial burden over time, this was done using the da_tracker workflow.^48^ For the quantification of LC3 volume over time, the bacterial burden section of the workflow was adapted to measure volume of signal in the LC3 channel instead of the bacterial channel.

For single MCV fluorescence quantification at defined time points, the workflow is illustrated in Figure S1 and ran in PyimageJ adapted from the da_tracker workflow. The source code is provided in supplementary material. In more details, the cells were segmented using Cellpose on the T-PMT channel. The bacteria channel is then isolated and the extracellular signal as well as signal in dead cells were removed. The bacterial centroid is then detected and collected using the plugin Trackmate.^86^ The bacteria were also detected using Fiji particle detection function and the generated ROIs were used to measure the centroid coordinates. Then, the coordinates of the centroids collected from Trackmate were compared to the coordinates of the ROIs centroid collected from Fiji using the nearest neighbor function from geopandas library in python. The refined ROI set representing was then used to measure the mean fluorescence intensity of the channels of interest. The code for this single MCV workflow is provided as supplementary material. For the determination of the relative bacterial burden at these defined time points, the bacterial fluorescence was analyzed to calculate the bacterial volume per cell, using the BBQ method described elsewhere. ^87^

### Data representation and statistics

Data were plotted using Graphpad Prism 9 or 10 except for the density map in Figure 5E, which was created using R. The figures were designed using Powerpoint and Inkscape, and the model design was made using Inkscape. All statistical tests were performed using Prism or excel. The correlation between two variables was tested using Pearson’s correlation test. Statistical differences between two groups were tested using paired Student *t*-tests or more than two groups using a one-way ANOVA followed by Dunnett *post-hoc* statistical test. The difference between two entire curves from time-lapses was done using the modified Chi-squared method on an Excel spreadsheet provided here.^88^ Comparisons with a *p-*value greater than 0.05 were considered significant.

## Data and code availability

The source code for the pyimagej workflow for single MCV quantification is provided as supplementary material. The movies referenced in this article were deposited here: https://data.mendeley.com/datasets/cvk8wnfm36/

Or can be obtained using the download links below : [Movie S1](https://data.mendeley.com/public-files/datasets/cvk8wnfm36/files/894b0b99-6c88-464f-8900-37fe1dcc153b/file_downloaded), [Movie S2](https://data.mendeley.com/public-files/datasets/cvk8wnfm36/files/4411d296-6a72-42d6-8810-372eb9af3fff/file_downloaded), [Movie S3](https://data.mendeley.com/public-files/datasets/cvk8wnfm36/files/36346594-525d-4f8e-94f0-a2f76fb24425/file_downloaded), [Movie S4](https://data.mendeley.com/public-files/datasets/cvk8wnfm36/files/0abd4cf1-beab-4011-903e-c5c6a12b2a0d/file_downloaded), [Movie S5](https://data.mendeley.com/public-files/datasets/cvk8wnfm36/files/5dca5a5b-e309-44a1-b635-0a4e299deca0/file_downloaded), [Movie S6](https://data.mendeley.com/public-files/datasets/cvk8wnfm36/files/2d33bede-0fae-4c5d-98ec-c79a675b8e43/file_downloaded), [Movie S7](https://data.mendeley.com/public-files/datasets/cvk8wnfm36/files/eae2a1a8-937c-4279-b079-c228d43006f8/file_downloaded), [Movie S8](https://data.mendeley.com/public-files/datasets/cvk8wnfm36/files/5e66d879-3fe7-4586-9953-5cc8154272d7/file_downloaded), [Movie S9](https://data.mendeley.com/public-files/datasets/cvk8wnfm36/files/3c784647-1ebd-4ac1-a633-b609c4a13d79/file_downloaded), [Movie S10](https://data.mendeley.com/public-files/datasets/cvk8wnfm36/files/b1408ed9-5b50-46a2-b35b-dbaa3d8b5410/file_downloaded), [Movie S11](https://data.mendeley.com/public-files/datasets/cvk8wnfm36/files/4b9dbffb-5e9e-49b1-8be8-69d50ccca5c5/file_downloaded), [Movie S12](https://data.mendeley.com/public-files/datasets/cvk8wnfm36/files/b947b43e-c6f2-4cdc-89f0-d5a328fd810a/file_downloaded), [Movie S13](https://data.mendeley.com/public-files/datasets/cvk8wnfm36/files/81eef5e4-19e5-47e9-9fe2-c8a9276f5a38/file_downloaded).

## Acknowledgements

We thank Dr Adrien Schahl for his insights in the optimization of the cell tracking calculation, and Dr Guillaume Ferré for his feedback on data analysis.

This work was supported by the National Institute of Allergy and Infectious Diseases (Grant R01AI139492 to V.B.). Purchase of the Zeiss LSM 980 Airyscan 2 was supported by Award Number 1S10OD025223-01A1 from the National Institute of Health.

Conceptualization: J.A., V.B.; Methodology: J.A., C.N.S.A., A.P., V.B..; Software: J.A., A.P.; Validation: J.A., V.B.; Formal analysis: J.A., A.T.P., A.P.; Investigation: J.A., A.T.P.; Resources: J.A., C.N.S.A., A.P., L.S.; Data curation: J.A.; Writing - original draft: J.A.; Writing - review & editing: J.A., C.N.S.A., V.B.; Visualization: J.A.; Supervision: J.A., V.B.; Project administration: J.A., V.B.; Funding acquisition: V.B.

We declare no conflicts of interest.

---

**Figure S1:**
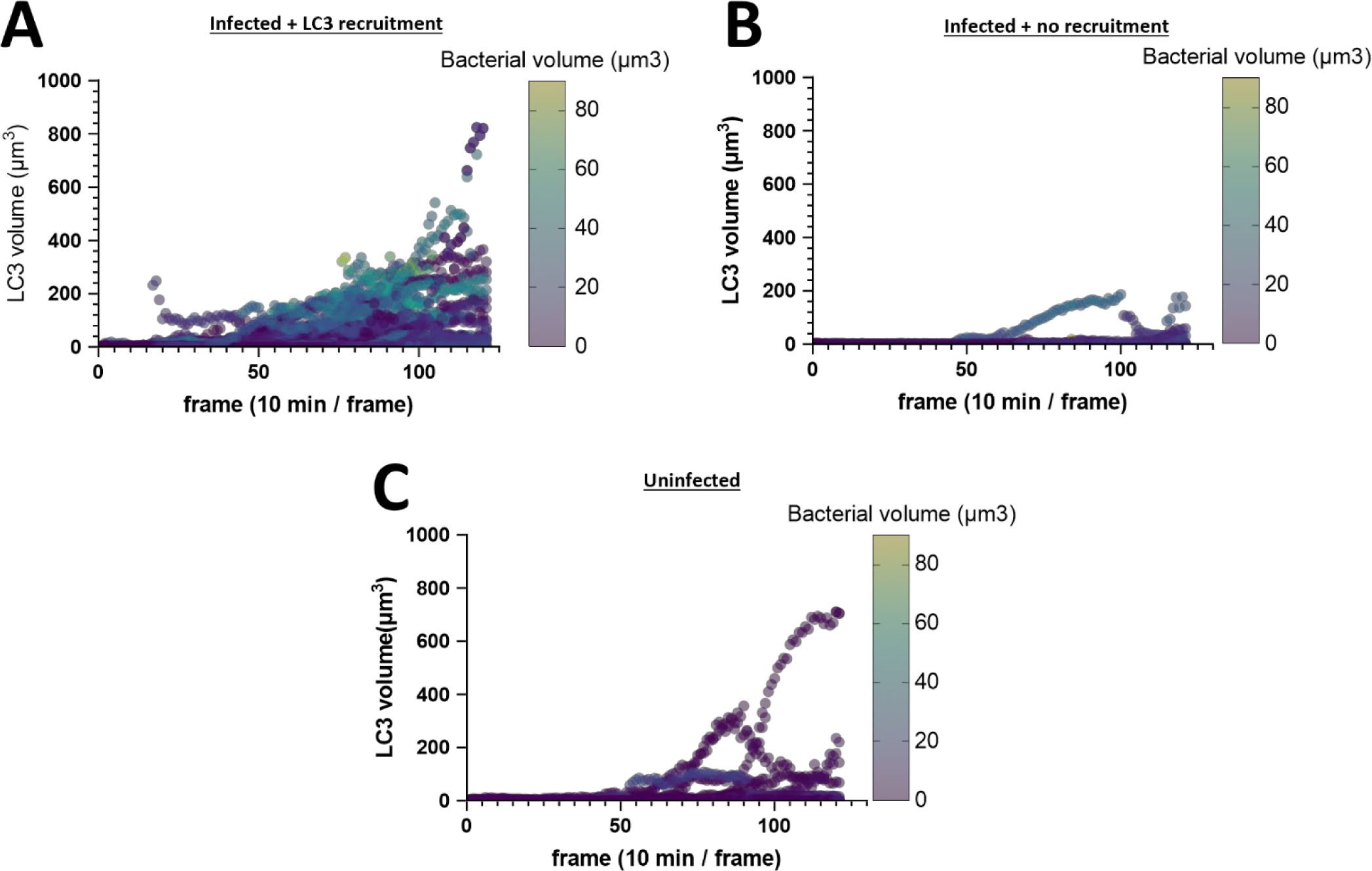
Quantification of autophagy induction over time. Single cell tracking and quantification of LC3 volume over time plotted on a 3 variables graph with the bacterial volume as a color dimension. Each dot represents the measurement in 1 cell in 1 frame. (A) Result of the quantification in cells exhibiting at least one temporary LC3 recruitment on at least one MCV (N_cells_ = 41, N_points_ = 4299). (B) Quantification in cells that don’t exhibit LC3 recruitment (N_cells_ = 11, N_points_ = 1222). (C) Quantification in uninfected cells (N_cells_ = 32, N_points_ = 3349).

**Figure S2:**
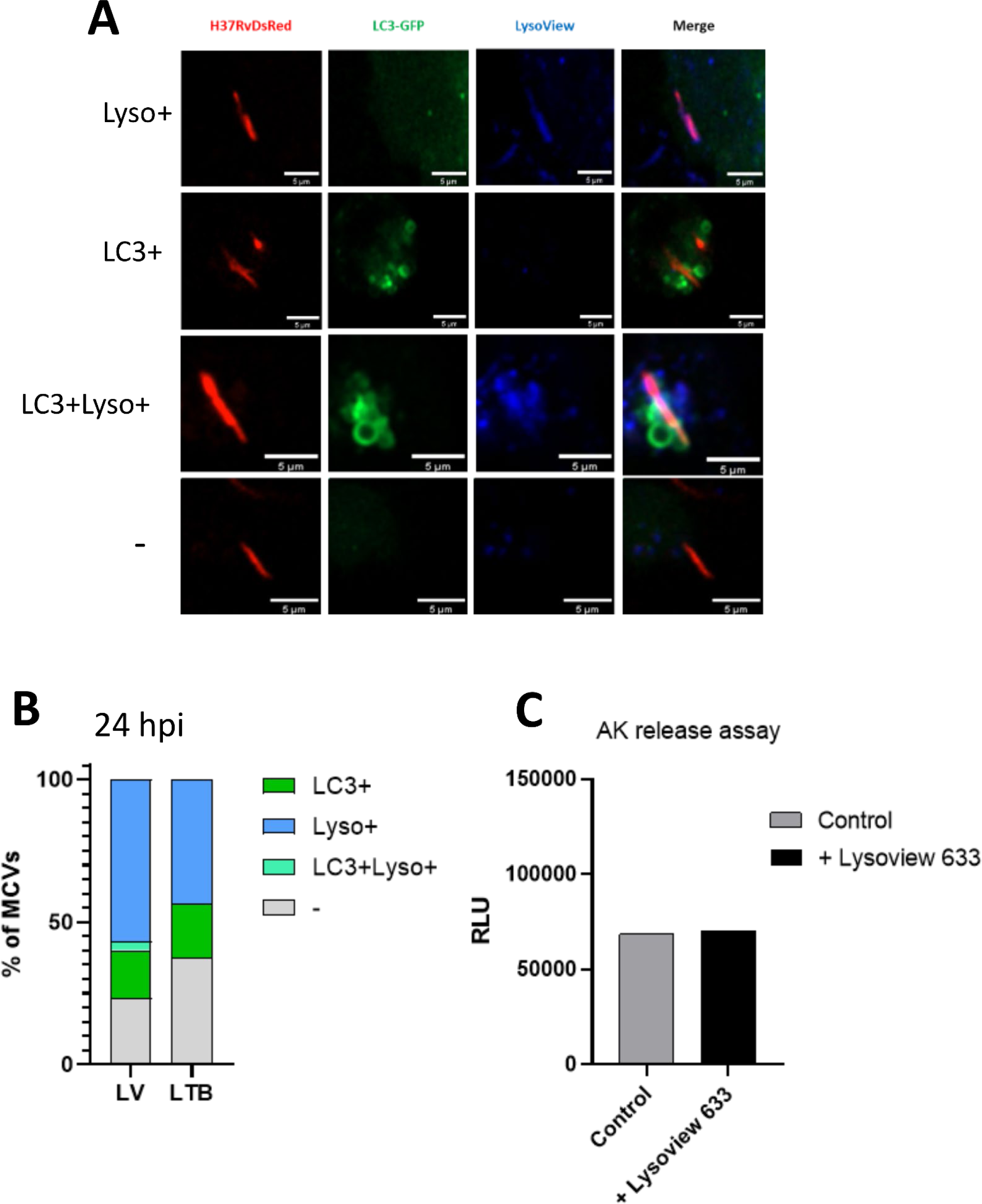
Comparison between Lysoview-633 and Lysotracker blue staining on infected cells. (A) Representative images of MCVs in THP-1-LC3-GFP cells negative (-), positive for Lysoview 633 (Lyso+), for LC3 (LC3+), and double positive (LC3+Lyso+). (B) quantification of the fraction of MCVs in THP-1 cells stained for lysoview 633 or lysotracker blue. (C) The cell death level was quantified by the adenylate Kinase (AK) release assay on THP-1-LC3-GFP treated or not with Lysoview 633 1:2000 for 48h. RLU = relative luminescence unit

**Figure S3.**
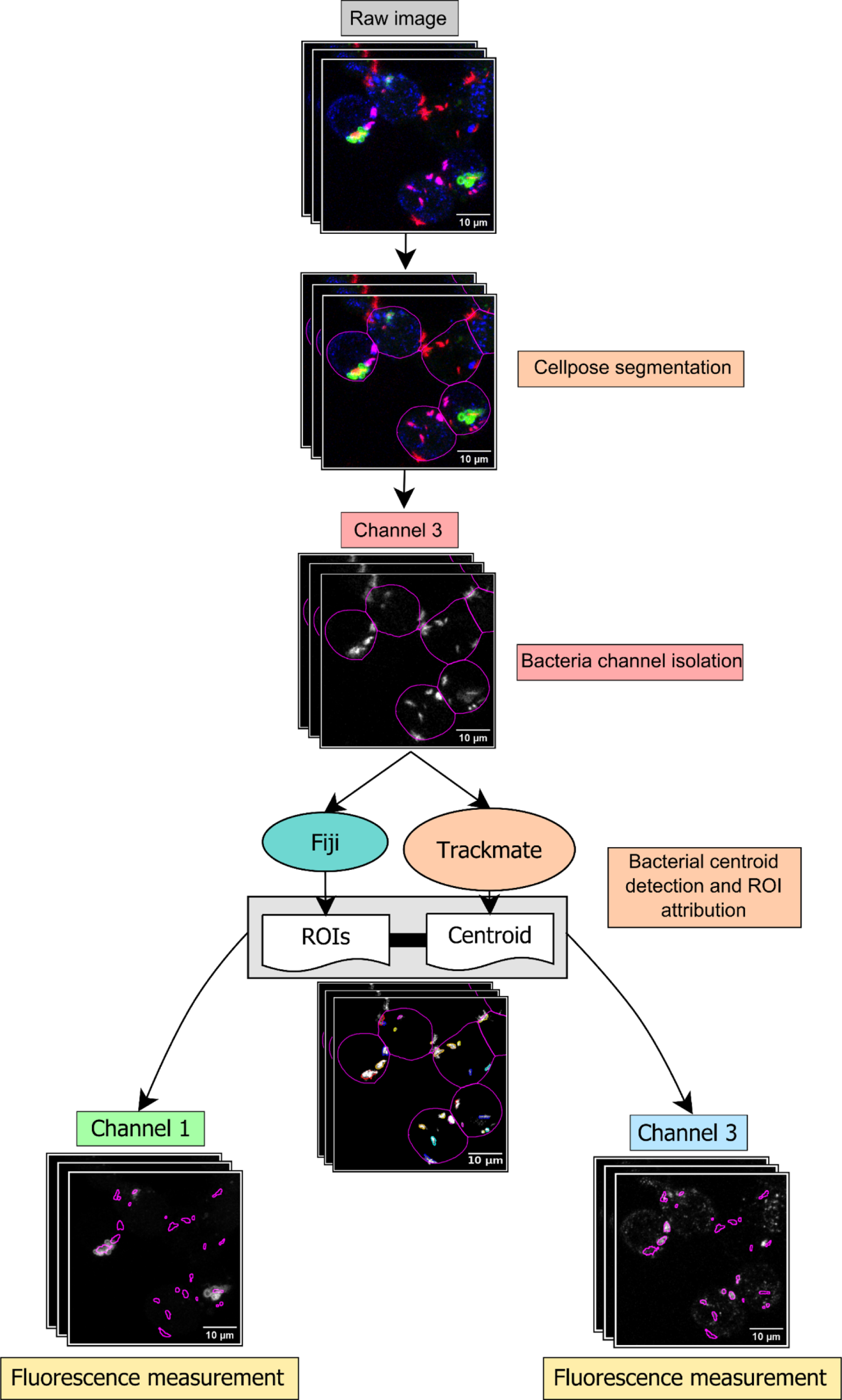
: Diagram of the image analysis workflow implemented for the fluorescence quantification on single MCV. The cells are first segmented using Cellpose. The ROIs obtained are applied on the bacterial channel to retain the signal only in living cells. The bacterial signal is then segmented on Fiji to create ROIs and measure the centroids coordinates, and analyzed using Trackmate to obtain the coodinates of the MCVs’ centroids. Their distance to the ROIs centroids is calculated to retain the closest using the nearest neighbor calculation. The selected ROIs were finally applied on the channels of interest and the mean fluorescence intensity is finally measured.

**Figure S4:**
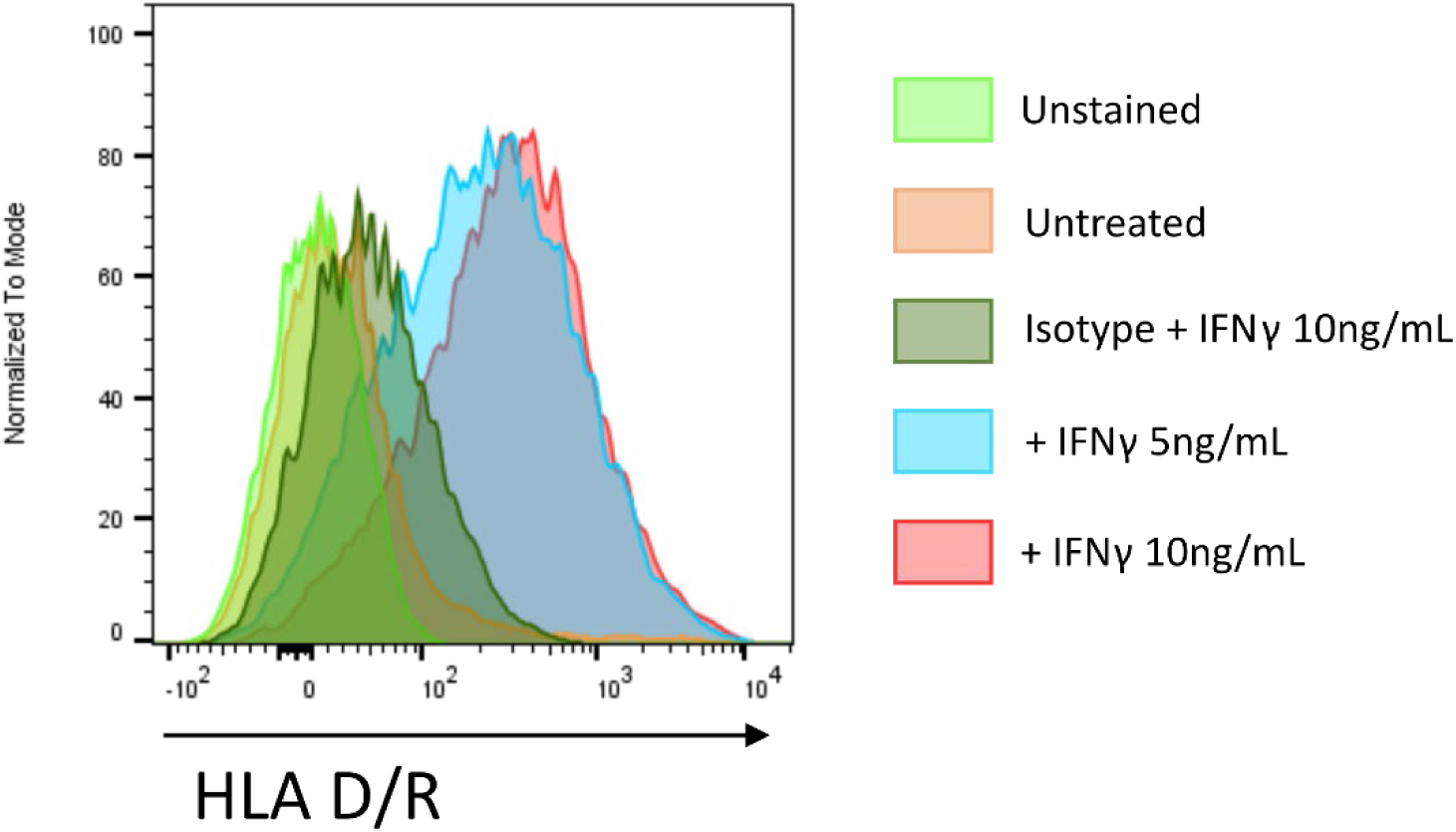
Effect of IFN-γ stimulation on MHC-II expression. THP-1 cells were treated with IFN-γ 5ng/mL, 10ng/mL overnight or left untreated. Cells were fixed and stained for MHC-II expression using HLA/DR antibody (BD). Some Treated cells with 10ng/mL were stained with isotype control. The staining was analyzed by flow cytometry.

**Figure S5:**
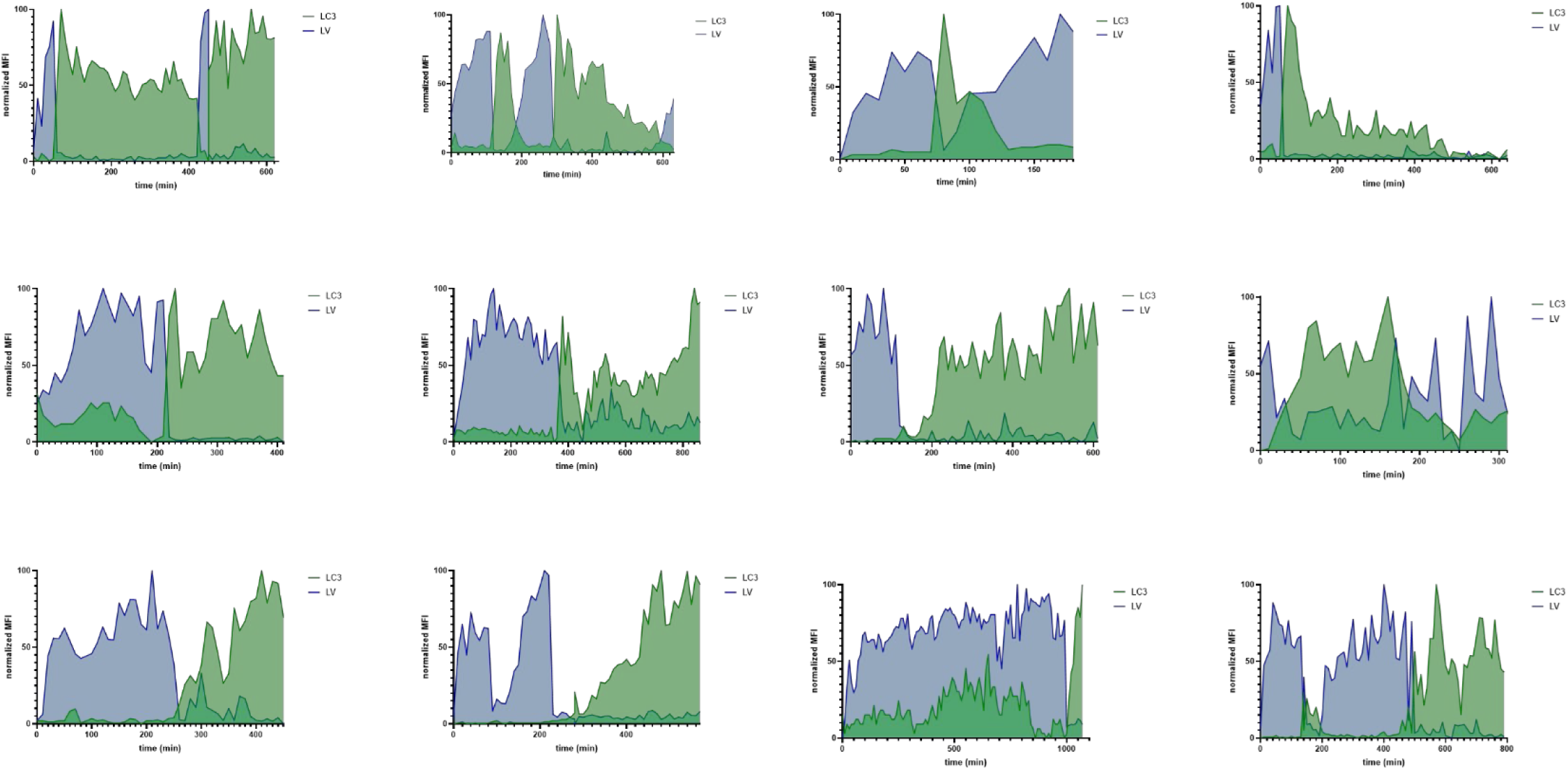
Fluorescence traces of LC3 and lysoview of individual MCVs. Compilation of the LC3 and lysoview fluorescence quantification on individual MCVs isolated from 4 independent experiments. Only the MCV exhibiting LC3 recruitment for more than 3h were retained.

**Figure S6:**
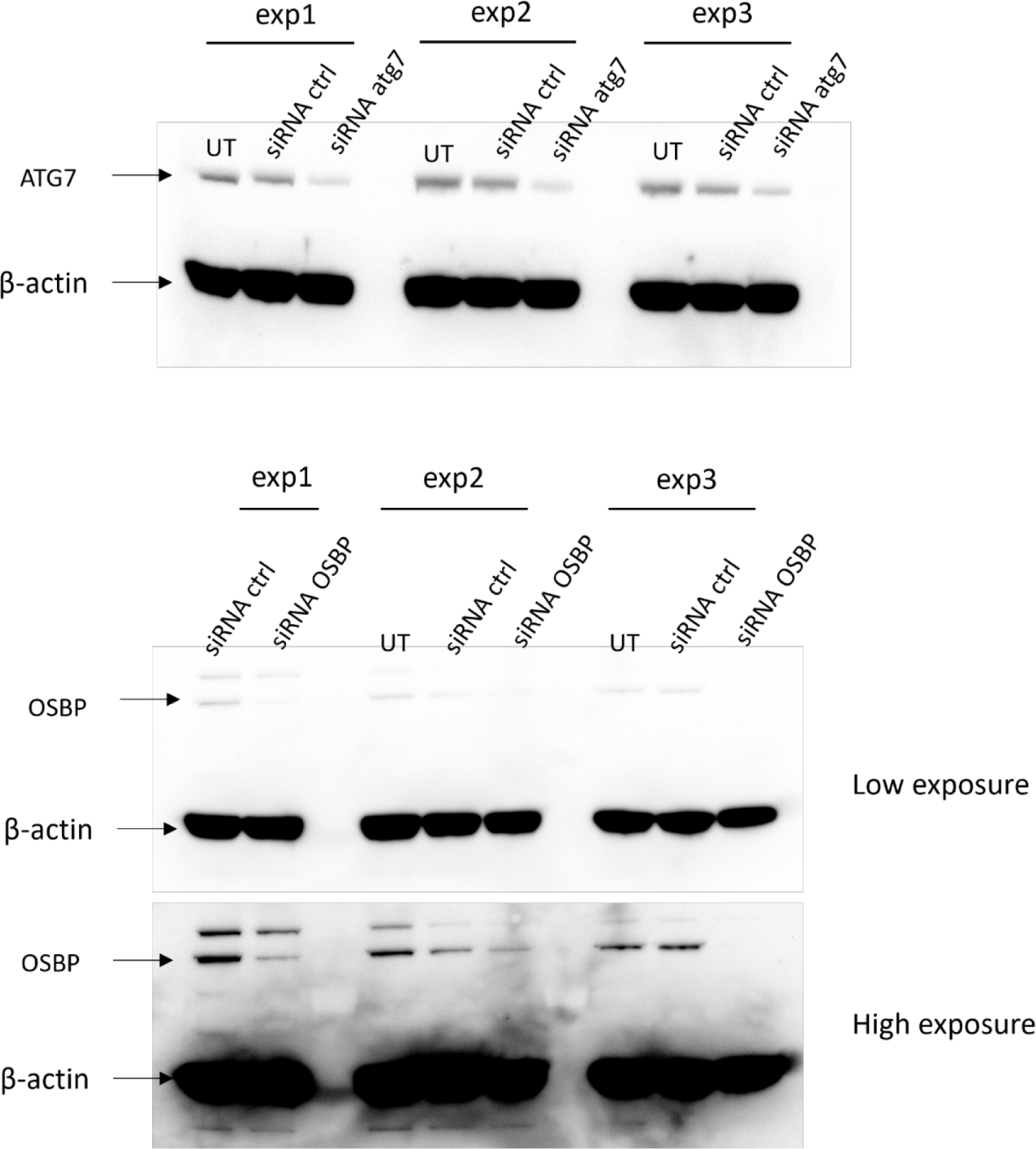
Relative abundance of ATG7 or OSBP in the THP-1-LC3-GFP cells after knock-down with siRNA. The cells were lysed and the lysate was analyzed by SDS-PAGE followed by western blot. The β-actin blotting served as a loading control. (top) Western blotting results from cells that were left untreated, treated with negative control siRNA, and siRNA against ATG7. The lysates correspond to the cells that were used for infection in Figure 6C-D. (Bottom) Western blotting results from cells untreated, treated with negative control siRNA, and siRNA against OSBP. The lysates correspond to the cells that were used for infection in Figure 6E-F. A low exposure and a high exposure are shown for clarity in the relative abundance of OSBP compared to the β-actin loading control.

**Figure S7:**
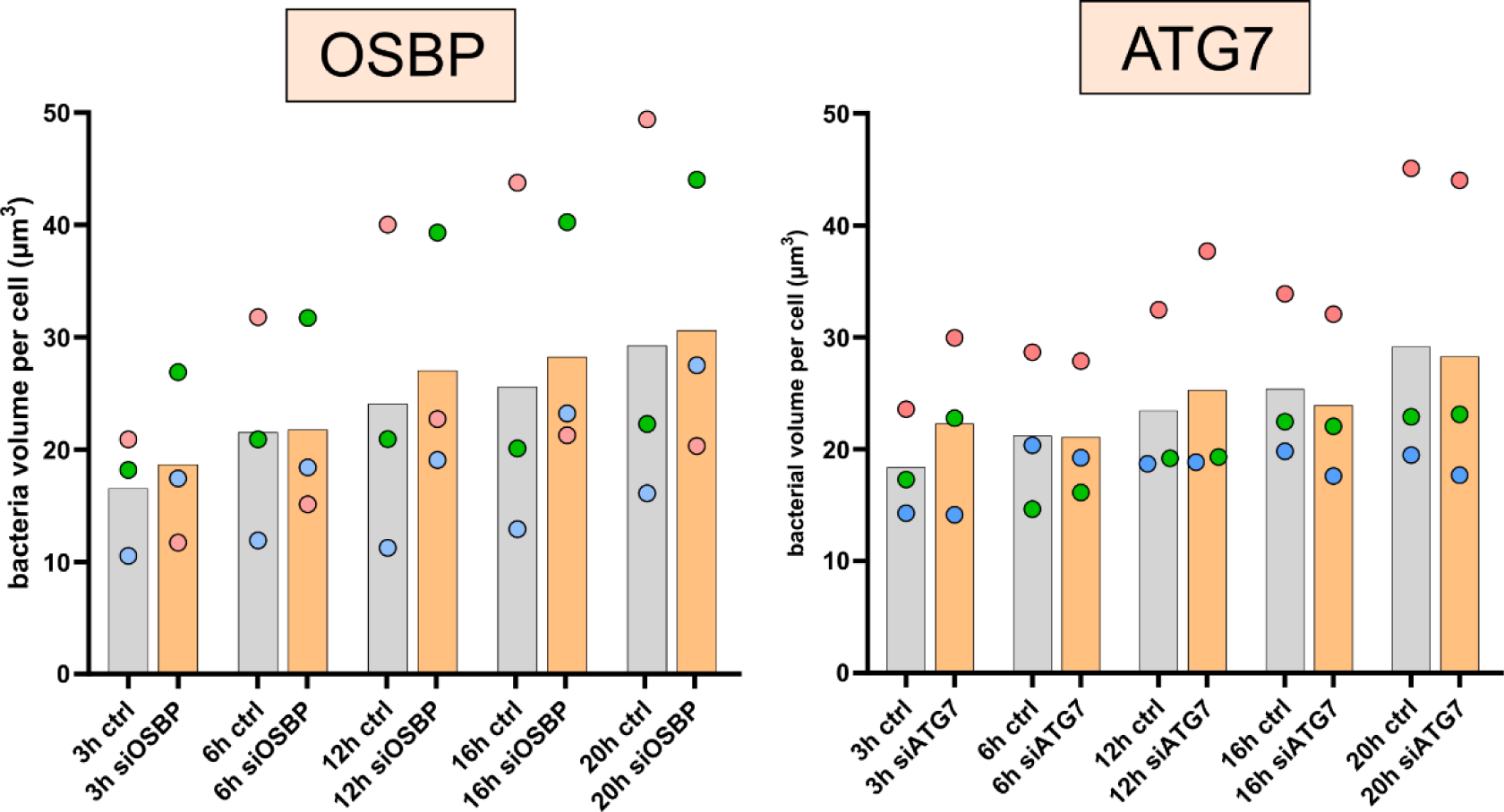
Evolution of relative bacterial volume per cell over time in OSBP and ATG7 knock-down cells. THP-1-LC3-GFP cells were transfected with siRNA against OSBP (left) or ATG7 (right), infected with Mtb, stained by Lysoview-633 and imaged by time-lapse confocal microscopy. The relative bacterial burden was determined by calculation of the bacterial volume per cell (see methods). These results corresponds to the raw values presented in the Figure 6D,F. The dots correspond to the average bacterial volume per cell and the color corresponds to the different independent experiments.

## Movie legends

**Movie S1: Representative field of view from a time lapse experiment monitoring LC3 recruitment to MCVs.**

THP-1 cells expressing LC3-GFP (green) were infected with H37Rv-DsRed (red) at an MOI 2 and imaged for 20h. A 10μm z-stack was acquired every 10 minutes. Time stamp format is hh:mm, and only one slice is shown. Example images are shown in Figure 1A.

**Movie S2: Representative time lapse of LC3 recruitment to the MCV followed by the exclusion of LC3 signal.**

A THP-1 cell expressing LC3-GFP (green) infected with H37Rv-DsRed (red) is shown. Time stamp format is hh:mm. Time 00:00 is the frame in which the bacteria visually enter the cell. Example images are shown in Figure 1C.

**Movie S3: Representative time lapse of LC3 recruitment to the MCV followed by the egress of the bacteria away from the LC3 vesicles.**

A THP-1 cell expressing LC3-GFP (green) infected with H37Rv-DsRed (red) is shown. Time stamp format is hh:mm. Time 00:00 is the frame in which the bacteria visually enter the cell. Example images are shown in Figure 1D.

**Movie S4: Representative time lapse of a LC3 recruitment to the MCV without sign of escape from the bacteria.**

A THP-1 cell expressing LC3-GFP (green) infected with H37Rv-DsRed (red) is shown. Time stamp format is hh:mm. Time 00:00 is the frame in which the bacteria visually enter the cell. Example images are shown in Figure 1E.

**Movie S5: Representative time lapse of an early LC3 recruitment to the MCV.**

A THP-1 cell expressing LC3-GFP (green) infected with H37Rv-DsRed (red) is shown. Time stamp format is hh:mm. Time 00:00 is the frame in which the bacteria visually enter the cell. The yellow outline is an ROI generated for the tracking of the MCV and fluorescence quantification. Example images are shown in Figure 2B.

**Movie 6: Representative time lapse of a late LC3 recruitment to the MCV.**

A THP-1 cell expressing LC3-GFP (green) infected with H37Rv-DsRed (red) is shown. Time stamp format is hh:mm. Time 00:00 is the frame in which the bacteria visually enter the cell. The yellow outline is an ROI generated for the tracking of the MCV and fluorescence quantification. Example images are shown in Figure 2D.

**Movie S7: Representative time lapse of multiple LC3 recruitments to the MCV.**

A THP-1 cell expressing LC3-GFP (green) infected with H37Rv-DsRed (red) is shown. Time stamp format is hh:mm. Time 00:00 is the frame in which the bacteria visually enter the cell. The yellow outline is an ROI generated for the tracking of the MCV and fluorescence quantification. Example images are shown in Figure 2F.

**Movie S8: Representative field of view from a time lapse experiment showing the specific tracking of cells that get infected and exhibit LC3 recruitment to the MCV.**

THP-1 cells expressing LC3-GFP (green) were infected with H37Rv-DsRed (red) at an MOI 2 and imaged for 20h. A 10μm z-stack was acquired every 10 minutes. Time stamp format is hh:mm, and only one slice is shown. Example images with or without the overlay are shown in Figure 3C.

**Movie S9: Representative field of view from a time lapse experiment showing the infection with Mtb of unstimulated cells with Mtb.**

THP-1 cells expressing LC3-GFP were infected with H37Rv-DsRed (red) at an MOI 2 and imaged for 20h. A 10μm z-stack was acquired every 10 minutes. Time stamp format is hh:mm, and only one slice is shown. Example images with or without the overlay are shown in Figure 4C top panel.

**Movie S10: Representative field of view from a time lapse experiment showing the infection with Mtb of cells pre-stimulated with IFN-γ.**

THP-1 cells expressing LC3-GFP were infected with H37Rv-DsRed (red) at an MOI 2 and imaged for 20h. A 10μm z-stack was acquired every 10 minutes. Time stamp format is hh:mm, and only one slice is shown. Example images with or without the overlay are shown in Figure 4C bottom panel.

**Movie S11: Representative time lapse of showing the staining of an MCV with lysoview or LC3 recruitment**

A THP-1 cell expressing LC3-GFP (green) stained with lysoview-633 (Blue) infected with H37Rv-DsRed (red) is shown. Time stamp format is hh:mm. Time 00:00 is the frame in which the bacteria visually enter the cell. The yellow outline is an ROI generated for the tracking of the MCV and fluorescence quantification. Example images are shown in Figure 5A.

**Movie S12: Representative time lapse of showing the staining of an MCV with lysoview or LC3 recruitment**

A THP-1 cell expressing LC3-GFP (green) stained with lysoview-633 (Blue) infected with H37Rv-DsRed (red) is shown. Time stamp format is hh:mm. Time 00:00 is the frame in which the bacteria visually enter the cell. The yellow outline is an ROI generated for the tracking of the MCV and fluorescence quantification. Example images are shown in Figure 5B.

**Movie S13: Representative time lapse showing the staining of an MCV with lysoview without LC3 recruitment**

A THP-1 cell expressing LC3-GFP (green) stained with lysoview-633 (Blue) infected with H37Rv-DsRed (red) is shown. The transmitted light channel is also displayed (gray). Time stamp format is hh:mm. Time 00:00 is the start of acquisition.

## References

1. Davis, J. M. & Ramakrishnan, L. The Role of the Granuloma in Expansion and Dissemination of Early Tuberculous Infection. Cell 136, 37–49 (2009).

2. Cohen, S. B. et al. Alveolar Macrophages Provide an Early Mycobacterium tuberculosis Niche and Initiate Dissemination. Cell Host Microbe 24, 439–446.e4 (2018).

3. Mahamed, D. et al. Intracellular growth of Mycobacterium tuberculosis after macrophage cell death leads to serial killing of host cells. eLife 6, e22028 (2017).

4. Lerner, T. R. et al. Mycobacterium tuberculosis replicates within necrotic human macrophages. J. Cell Biol. 216, 583–594 (2017).

5. Queval, C. J., Brosch, R. & Simeone, R. The Macrophage: A Disputed Fortress in the Battle against Mycobacterium tuberculosis. Front. Microbiol. 8, 2284 (2017).

6. Augenstreich, J. & Briken, V. Host Cell Targets of Released Lipid and Secreted Protein Effectors of Mycobacterium tuberculosis. Front. Cell. Infect. Microbiol. 10, (2020).

7. Chandra, P., Grigsby, S. J. & Philips, J. A. Immune evasion and provocation by Mycobacterium tuberculosis. Nat. Rev. Microbiol. 1–17 (2022) doi:10.1038/s41579-022-00763-4.

8. Deretic, V. Autophagy in inflammation, infection, and immunometabolism. Immunity 54, 437–453 (2021).

9. Upadhyay, S. & Philips, J. A. LC3-associated phagocytosis: host defense and microbial response. Curr. Opin. Immunol. 60, 81–90 (2019).

10. Gutierrez, M. G. et al. Autophagy Is a Defense Mechanism Inhibiting BCG and Mycobacterium tuberculosis Survival in Infected Macrophages. Cell 119, 753–766 (2004).

11. Manzanillo, P. S. et al. The ubiquitin ligase parkin mediates resistance to intracellular pathogens. Nature 501, 512–516 (2013).

12. Aylan, B. et al. ATG7 and ATG14 restrict cytosolic and phagosomal Mycobacterium tuberculosis replication in human macrophages. Nat. Microbiol. 1–16 (2023) doi:10.1038/s41564-023-01335-9.

13. Kinsella, R. L. et al. Autophagy prevents early proinflammatory responses and neutrophil recruitment during Mycobacterium tuberculosis infection without affecting pathogen burden in macrophages. PLOS Biol. 21, e3002159 (2023).

14. Feng, S. et al. Autophagy promotes efficient T cell responses to restrict high-dose Mycobacterium tuberculosis infection in mice. Nat. Microbiol. 9, 684–697 (2024).

15. Köster, S., et al. *Mycobacterium tuberculosis* is protected from NADPH oxidase and LC3-associated phagocytosis by the LCP protein CpsA. Proc. Natl. Acad. Sci. 114, E8711–E8720 (2017).

16. Bell, S. L., Lopez, K. L., Cox, J. S., Patrick, K. L. & Watson, R. O. Galectin-8 Senses Phagosomal Damage and Recruits Selective Autophagy Adapter TAX1BP1 To Control Mycobacterium tuberculosis Infection in Macrophages. mBio 12, e01871–20 (2021).

17. Watson, R. O., Manzanillo, P. S. & Cox, J. S. Extracellular M. tuberculosis DNA Targets Bacteria for Autophagy by Activating the Host DNA-Sensing Pathway. Cell 150, 803–815 (2012).

18. Watson, R. O. et al. The Cytosolic Sensor cGAS Detects Mycobacterium tuberculosis DNA to Induce Type I Interferons and Activate Autophagy. Cell Host Microbe 17, 811–819 (2015).

19. Franco, L. H. et al. The Ubiquitin Ligase Smurf1 Functions in Selective Autophagy of Mycobacterium tuberculosis and Anti-tuberculous Host Defense. Cell Host Microbe 21, 59–72 (2017).

20. Sanjuan, M. A. et al. Toll-like receptor signalling in macrophages links the autophagy pathway to phagocytosis. Nature 450, 1253–1257 (2007).

21. Huang, J. et al. Activation of antibacterial autophagy by NADPH oxidases. Proc. Natl. Acad. Sci. 106, 6226–6231 (2009).

22. Vandal, O. H., Nathan, C. F. & Ehrt, S. Acid Resistance in Mycobacterium tuberculosis. J. Bacteriol. 191, 4714–4721 (2009).

23. Cemma, M., Grinstein, S. & Brumell, J. H. Autophagy proteins are not universally required for phagosome maturation. Autophagy 12, 1440–1446 (2016).

24. Cardenal-Muñoz, E. et al. Mycobacterium marinum antagonistically induces an autophagic response while repressing the autophagic flux in a TORC1- and ESX-1-dependent manner. PLOS Pathog. 13, e1006344 (2017).

25. López-Jiménez, A. T. et al. The ESCRT and autophagy machineries cooperate to repair ESX-1-dependent damage at the Mycobacterium-containing vacuole but have opposite impact on containing the infection. PLOS Pathog. 14, e1007501 (2018).

26. Romagnoli, A. et al. ESX-1 dependent impairment of autophagic flux by Mycobacterium tuberculosis in human dendritic cells. Autophagy 8, 1357–1370 (2012).

27. Chandra, P. et al. Mycobacterium tuberculosis Inhibits RAB7 Recruitment to Selectively Modulate Autophagy Flux in Macrophages. Sci. Rep. 5, 16320 (2015).

28. Bah, A. et al. The Lipid Virulence Factors of Mycobacterium tuberculosis Exert Multilayered Control over Autophagy-Related Pathways in Infected Human Macrophages. Cells 9, 666 (2020).

29. Saini, N. K. et al. Suppression of autophagy and antigen presentation by Mycobacterium tuberculosis PE_PGRS47. Nat. Microbiol. 1, 1–12 (2016).

30. Shin, D.-M. et al. Mycobacterium tuberculosis Eis Regulates Autophagy, Inflammation, and Cell Death through Redox-dependent Signaling. PLOS Pathog. 6, e1001230 (2010).

31. Strong, E. J. et al. Identification of Autophagy-Inhibiting Factors of Mycobacterium tuberculosis by High-Throughput Loss-of-Function Screening. Infect. Immun. 88, e00269–20 (2020).

32. Strong, E. J., Ng, T. W., Porcelli, S. A. & Lee, S. Mycobacterium tuberculosis PE_PGRS20 and PE_PGRS47 Proteins Inhibit Autophagy by Interaction with Rab1A. mSphere 6, e00549–21 (2021).

33. Bernard, E. M. et al. M. tuberculosis infection of human iPSC-derived macrophages reveals complex membrane dynamics during xenophagy evasion. J. Cell Sci. 134, jcs252973 (2020).

34. Armstrong, J. A. & Hart, P. D. Phagosome-lysosome interactions in cultured macrophages infected with virulent tubercle bacilli. Reversal of the usual nonfusion pattern and observations on bacterial survival. J. Exp. Med. 142, 1–16 (1975).

35. Sturgill-Koszycki, S. et al. Lack of Acidification in Mycobacterium Phagosomes Produced by Exclusion of the Vesicular Proton-ATPase. Science 263, 678–681 (1994).

36. Clemens, D. L. & Horwitz, M. A. Characterization of the Mycobacterium tuberculosis phagosome and evidence that phagosomal maturation is inhibited. J. Exp. Med. 181, 257–270 (1995).

37. Hasan, Z. et al. Isolation and characterization of the mycobacterial phagosome: segregation from the endosomal/lysosomal pathway. Mol. Microbiol. 24, 545–553 (1997).

38. Fratti, R. A., Backer, J. M., Gruenberg, J., Corvera, S. & Deretic, V. Role of phosphatidylinositol 3-kinase and Rab5 effectors in phagosomal biogenesis and mycobacterial phagosome maturation arrest. J. Cell Biol. 154, 631–644 (2001).

39. Bach, H., Papavinasasundaram, K. G., Wong, D., Hmama, Z. & Av-Gay, Y. Mycobacterium tuberculosis Virulence Is Mediated by PtpA Dephosphorylation of Human Vacuolar Protein Sorting 33B. Cell Host Microbe 3, 316–322 (2008).

40. van der Wel, N. et al. M. tuberculosis and M. leprae Translocate from the Phagolysosome to the Cytosol in Myeloid Cells. Cell 129, 1287–1298 (2007).

41. Simeone, R. et al. Phagosomal Rupture by Mycobacterium tuberculosis Results in Toxicity and Host Cell Death. PLoS Pathog. 8, e1002507 (2012).

42. Levitte, S. et al. Mycobacterial Acid Tolerance Enables Phagolysosomal Survival and Establishment of Tuberculous Infection In Vivo. Cell Host Microbe 20, 250–258 (2016).

43. Gutierrez, M. G. & Enninga, J. Intracellular niche switching as host subversion strategy of bacterial pathogens. Curr. Opin. Cell Biol. 76, 102081 (2022).

44. Bussi, C. & Gutierrez, M. G. Mycobacterium tuberculosis infection of host cells in space and time. FEMS Microbiol. Rev. 43, 341–361 (2019).

45. Beckwith, K. S. et al. Plasma membrane damage causes NLRP3 activation and pyroptosis during Mycobacterium tuberculosis infection. Nat. Commun. 11, 2270 (2020).

46. Rutschmann, O., Toniolo, C. & McKinney, J. D. Preexisting Heterogeneity of Inducible Nitric Oxide Synthase Expression Drives Differential Growth of Mycobacterium tuberculosis in Macrophages. mBio 0, e02251–22 (2022).

47. Toniolo, C., Dhar, N. & McKinney, J. D. Uptake-independent killing of macrophages by extracellular Mycobacterium tuberculosis aggregates. EMBO J. 42, e113490 (2023).

48. Augenstreich, J., Poddar, A., Belew, A. T., El-Sayed, N. M. & Briken, V. da_Tracker: Automated workflow for high throughput single cell and single phagosome tracking in infected cells. 2024.04.10.588863 Preprint at 10.1101/2024.04.10.588863 (2024).

49. Srinivasan, L. et al. Identification of a Transcription Factor That Regulates Host Cell Exit and Virulence of Mycobacterium tuberculosis. PLOS Pathog. 12, e1005652 (2016).

50. Gao, W. et al. Biochemical Isolation and Characterization of the Tubulovesicular LC3-positive Autophagosomal Compartment *. J. Biol. Chem. 285, 1371–1383 (2010).

51. Lerner, T. R., et al. *Mycobacterium tuberculosis* cords within lymphatic endothelial cells to evade host immunity. JCI Insight 5, (2020).

52. Wong, K.-W. & Jacobs Jr, W. R. Critical role for NLRP3 in necrotic death triggered by Mycobacterium tuberculosis: NLRP3 mediates M. tuberculosis-promoted necrosis. Cell. Microbiol. 13, 1371–1384 (2011).

53. Jia, J. et al. Galectin-3 Coordinates a Cellular System for Lysosomal Repair and Removal. Dev. Cell 52, 69–87.e8 (2020).

54. Anand, A. et al. ER-dependent membrane repair of mycobacteria-induced vacuole damage. mBio 14, e00943–23 (2023).

55. Repnik, U. et al. L-leucyl-L-leucine methyl ester does not release cysteine cathepsins to the cytosol but inactivates them in transiently permeabilized lysosomes. J. Cell Sci. 130, 3124–3140 (2017).

56. Radulovic, M. et al. ESCRT-mediated lysosome repair precedes lysophagy and promotes cell survival. EMBO J. 37, e99753 (2018).

57. Skowyra, M. L., Schlesinger, P. H., Naismith, T. V. & Hanson, P. I. Triggered recruitment of ESCRT machinery promotes endolysosomal repair. Science 360, eaar5078 (2018).

58. Radulovic, M. et al. Lysosome repair by ER-mediated cholesterol transfer. 2022.09.26.509457 Preprint at 10.1101/2022.09.26.509457 (2022).

59. Schnettger, L. et al. A Rab20-Dependent Membrane Trafficking Pathway Controls M. tuberculosis Replication by Regulating Phagosome Spaciousness and Integrity. Cell Host Microbe 21, 619–628.e5 (2017).

60. Kreibich, S. et al. Autophagy Proteins Promote Repair of Endosomal Membranes Damaged by the Salmonella Type Three Secretion System 1. Cell Host Microbe 18, 527–537 (2015).

61. Baker, J. J., Johnson, B. K. & Abramovitch, R. B. Slow growth of Mycobacterium tuberculosis at acidic pH is regulated by phoPR and host-associated carbon sources. Mol. Microbiol. 94, 56–69 (2014).

62. Frigui, W. et al. Control of M. tuberculosis ESAT-6 Secretion and Specific T Cell Recognition by PhoP. PLOS Pathog. 4, e33 (2008).

63. Ma, Y., Keil, V. & Sun, J. Characterization of *Mycobacterium tuberculosis* EsxA Membrane Insertion: ROLES OF N- AND C-TERMINAL FLEXIBLE ARMS AND CENTRAL HELIX-TURN-HELIX MOTIF. J. Biol. Chem. 290, 7314–7322 (2015).

64. De Leon, J., et al. *Mycobacterium tuberculosis* ESAT-6 Exhibits a Unique Membrane-interacting Activity That Is Not Found in Its Ortholog from Non-pathogenic *Mycobacterium smegmatis*. J. Biol. Chem. 287, 44184–44191 (2012).

65. de Jonge, M. I. et al. ESAT-6 from Mycobacterium tuberculosis Dissociates from Its Putative Chaperone CFP-10 under Acidic Conditions and Exhibits Membrane-Lysing Activity. J. Bacteriol. 189, 6028–6034 (2007).

66. Simeone, R. et al. Cytosolic Access of Mycobacterium tuberculosis: Critical Impact of Phagosomal Acidification Control and Demonstration of Occurrence In Vivo. PLOS Pathog. 11, e1004650 (2015).

67. Augenstreich, J. et al. ESX-1 and phthiocerol dimycocerosates of *Mycobacterium tuberculosis* act in concert to cause phagosomal rupture and host cell apoptosis. Cell. Microbiol. 19, e12726 (2017).

68. Pethe, K. et al. Isolation of Mycobacterium tuberculosis mutants defective in the arrest of phagosome maturation. Proc. Natl. Acad. Sci. 101, 13642–13647 (2004).

69. Jutras, I. et al. Modulation of the Phagosome Proteome by Interferon-γ *. Mol. Cell. Proteomics 7, 697–715 (2008).

70. Wong, K.-W. & Jacobs, W. R. Mycobacterium tuberculosis exploits human interferon γ to stimulate macrophage extracellular trap formation and necrosis. J. Infect. Dis. 208, 109–119 (2013).

71. Simpson, D. S. et al. Interferon-γ primes macrophages for pathogen ligand-induced killing via a caspase-8 and mitochondrial cell death pathway. Immunity 55, 423–441.e9 (2022).

72. Golovkine, G. R. et al. Autophagy restricts Mycobacterium tuberculosis during acute infection in mice. Nat. Microbiol. 1–14 (2023) doi:10.1038/s41564-023-01354-6.

73. Kimmey, J. M. et al. Unique role for ATG5 in neutrophil-mediated immunopathology during M. tuberculosis infection. Nature 528, 565–569 (2015).

74. Deretic, V. Atg8ylation as a host-protective mechanism against Mycobacterium tuberculosis. Front. Tuberc. 1, (2023).

75. Peña-Martinez, C., Rickman, A. D. & Heckmann, B. L. Beyond autophagy: LC3-associated phagocytosis and endocytosis. Sci. Adv. 8, eabn1702 (2022).

76. Johnsson, K., Froland, W. A. & Schultz, P. G. Overexpression, Purification, and Characterization of the Catalase-peroxidase KatG from Mycobacterium tuberculosis*. J. Biol. Chem. 272, 2834–2840 (1997).

77. Braunstein, M., Espinosa, B. J., Chan, J., Belisle, J. T. & R. Jacobs Jr, W. SecA2 functions in the secretion of superoxide dismutase A and in the virulence of Mycobacterium tuberculosis. Mol. Microbiol. 48, 453–464 (2003).

78. Miller, J. L., Velmurugan, K., Cowan, M. J. & Briken, V. The Type I NADH Dehydrogenase of Mycobacterium tuberculosis Counters Phagosomal NOX2 Activity to Inhibit TNF-α-Mediated Host Cell Apoptosis. PLOS Pathog. 6, e1000864 (2010).

79. Srivastava, S., Battu, M. B., Khan, M. Z., Nandicoori, V. K. & Mukhopadhyay, S. Mycobacterium tuberculosis PPE2 Protein Interacts with p67phox and Inhibits Reactive Oxygen Species Production. J. Immunol. 203, 1218–1229 (2019).

80. Hooper, K. M. et al. V-ATPase is a universal regulator of LC3-associated phagocytosis and non-canonical autophagy. J. Cell Biol. 221, e202105112 (2022).

81. Biswas, V. K. et al. NCoR1 controls Mycobacterium tuberculosis growth in myeloid cells by regulating the AMPK-mTOR-TFEB axis. PLOS Biol. 21, e3002231 (2023).

82. Quigley, J. et al. The Cell Wall Lipid PDIM Contributes to Phagosomal Escape and Host Cell Exit of Mycobacterium tuberculosis. mBio 8, mBio.00148-17, e00148-17 (2017).

83. Schindelin, J., et al. Fiji: an open-source platform for biological-image analysis. Nat. Methods 9, 676–682 (2012).

84. Rueden, C. T. et al. PyImageJ: A library for integrating ImageJ and Python. Nat. Methods 1–2 (2022) doi:10.1038/s41592-022-01655-4.

85. Schnettger, L. & Gutierrez, M. G. Quantitative Spatiotemporal Analysis of Phagosome Maturation in Live Cells. in Phagocytosis and Phagosomes: Methods and Protocols (ed. Botelho, R.) 169–184 (Springer, New York, NY, 2017). doi:10.1007/978-1-4939-6581-6_11.

86. Ershov, D. et al. TrackMate 7: integrating state-of-the-art segmentation algorithms into tracking pipelines. Nat. Methods 19, 829–832 (2022).

87. Augenstreich, J. et al. BBQ methods: streamlined workflows for bacterial burden quantification in infected cells by confocal microscopy. Biol. Open 13, bio060189 (2024).

88. Hristova, K. & Wimley, W. C. Determining the statistical significance of the difference between arbitrary curves: A spreadsheet method. PLOS ONE 18, e0289619 (2023).

89. Stringer, C. & Pachitariu, M. Cellpose 2.0: how to train your own model. 2022.04.01.486764 Preprint at 10.1101/2022.04.01.486764 (2022).

