## Supplementary material for "Dynamic Interplay of Autophagy and Membrane Repair During *Mycobacterium tuberculosis* Infection": Source_code

### single_bact_quant

January 25, 2024

#### 1 Modules and libraries

[ ]:
```
from cellpose import models, io
from cellpose.io import *
import glob
import imagej
from jpype import JArray, JInt
import matplotlib.pyplot as plt
import os
import re
import pandas
from pandas import DataFrame
from pathlib import Path
import scyjava
import shutil
import tkinter as tk
from tkinter import filedialog
from PIL import Image
import sys
import os
```

#### 2 Fiji initialization

[ ]:
```
scyjava.config.add_option('-Xmx30g')
#ij = imagej.init('/home/saka/fiji-linux64/Fiji.app', mode = 'interactive')
ij = imagej.init('/home/saka/sw/local/fiji/2023', mode='interactive')
ij.ui().showUI()
ij.getVersion()
```

#### 3 Import of Fiji plugins and Trackmate settings

[ ]:
```
showPolygonRoi = scyjava.jimport('ij.gui.PolygonRoi')
Overlay = scyjava.jimport('ij.gui.Overlay')
Regions = scyjava.jimport('net.imglib2.roi.Regions')
LabelRegions = scyjava.jimport('net.imglib2.roi.labeling.LabelRegions')
```

```
ZProjector = scyjava.jimport('ij.plugin.ZProjector')()
Duplicator = scyjava.jimport('ij.plugin.Duplicator')()
ov = Overlay()
Model =  scyjava.jimport('fiji.plugin.trackmate.Model')
Settings= scyjava.jimport('fiji.plugin.trackmate.Settings')
TrackMate = scyjava.jimport('fiji.plugin.trackmate.TrackMate')
Settings= scyjava.jimport('fiji.plugin.trackmate.Settings')
TrackMate = scyjava.jimport('fiji.plugin.trackmate.TrackMate')
Logger= scyjava.jimport('fiji.plugin.trackmate.Logger')
DetectorKeys= scyjava.jimport('fiji.plugin.trackmate.detection.DetectorKeys')
ExportTracksToXML= scyjava.jimport('fiji.plugin.trackmate.action.
  ↪ExportTracksToXML')
TmXmlWriter= scyjava.jimport('fiji.plugin.trackmate.io.TmXmlWriter')
LogRecorder = scyjava.jimport('fiji.plugin.trackmate.util.LogRecorder')
SparseLAPTrackerFactory= scyjava.jimport('fiji.plugin.trackmate.tracking.
  ↪jaqaman.SparseLAPTrackerFactory')
TMUtils = scyjava.jimport('fiji.plugin.trackmate.util.TMUtils')
HyperStackDisplayer = scyjava.jimport('fiji.plugin.trackmate.visualization.
  ↪hyperstack.HyperStackDisplayer')
SelectionModel = scyjava.jimport('fiji.plugin.trackmate.SelectionModel')
CellposeDetectorFactory = scyjava.jimport('fiji.plugin.trackmate.cellpose.
  ↪CellposeDetectorFactory')
FeatureFilter = scyjava.jimport('fiji.plugin.trackmate.features.FeatureFilter')
DisplaySetting = scyjava.jimport('fiji.plugin.trackmate.gui.displaysettings.
  ↪DisplaySettings')
DisplaySettingsIO = scyjava.jimport('fiji.plugin.trackmate.gui.displaysettings.
  ↪DisplaySettingsIO')
CaptureOverlayAction = scyjava.jimport('fiji.plugin.trackmate.action.
  ↪CaptureOverlayAction')
PretrainedModel= scyjava.jimport('fiji.plugin.trackmate.cellpose.
  ↪CellposeSettings.PretrainedModel')
ThresholdDetectorFactory= scyjava.jimport('fiji.plugin.trackmate.detection.
  ↪ThresholdDetectorFactory')
TrackScheme = scyjava.jimport('fiji.plugin.trackmate.visualization.trackscheme.
  ↪TrackScheme')
TrackTableView = scyjava.jimport('fiji.plugin.trackmate.visualization.table.
  ↪TrackTableView')
AllSpotsTableView = scyjava.jimport('fiji.plugin.trackmate.visualization.table.
  ↪AllSpotsTableView')

rm = ij.RoiManager.getRoiManager()
```

### 4 Main directory choice and create of future sub-directories containing different outputs

[ ]:
```
#define your parent directory
base_path = '/home/saka/Documents/Lab_stuff/confocal/INFg/exp583/scene8'

# define time point of interest by its frame number
wanted_frame = 37

#Creation of different directory for outputs
directory_path = f"{base_path}/frame{wanted_frame}"
if not os.path.exists(directory_path):
    os.makedirs(directory_path)
frame_path = directory_path + "/frame"
if not os.path.exists(frame_path):
    os.makedirs(frame_path)
measurement_path = directory_path + "/measurement/"
if not os.path.exists(measurement_path):
    os.makedirs(measurement_path)
```

[ ]:
```
# open raw file
raw_image = ij.io().open('/home/saka/Documents/Lab_stuff/confocal/INFg/exp583/
 ↪scene8/Scene8.czi')
```

### 5 generation of single z multi-channel image and single channel z-stacks images

[ ]:
```
format = f'Tiff'
wanted_channel = 3
wanted_z = 3
image = raw_image[:, :, wanted_channel, wanted_z, wanted_frame]
frame = ij.py.to_imageplus(image)
frame.setDimensions(1, 1, 121)
#ij.ui().show(lc3)
result_path = f"{frame_path}/frame.tif"
ij.IJ.saveAs(frame, "Tiff", ij.py.to_java(result_path))
```

[ ]:
```
bact_channel = 1
channel = raw_image[:, :, bact_channel, :, wanted_frame]
bact = ij.py.to_imageplus(channel)
bact.setDimensions(1, 11, 1)
#ij.ui().show(bact)
result_path = f"{frame_path}/channel_bact.tif"
ij.IJ.saveAs(bact, "Tiff", ij.py.to_java(result_path))
```

[ ]:
```
format = f'Tiff'

LC3_channel = 0
channel = raw_image[:, :, LC3_channel, :, wanted_frame]
lc3 = ij.py.to_imageplus(channel)
lc3.setDimensions(1, 11, 1)
#ij.ui().show(lc3)
result_path = f"{directory_path}/LC3_channel.tif"
ij.IJ.saveAs(lc3, "Tiff", ij.py.to_java(result_path))

LV_channel = 2
channel = raw_image[:, :, LV_channel, :, wanted_frame]
lv = ij.py.to_imageplus(channel)
lv.setDimensions(1, 11, 1)
result_path = f"{directory_path}/LV_channel.tif"
ij.IJ.saveAs(lv, "Tiff", ij.py.to_java(result_path))
```

#### 6 cellpose segmentation on single z image - gray channel and collection of ROIs

[ ]:
```
image_cp = f"{frame_path}/frame.tif"
model = models.CellposeModel(gpu=True, model_type='CP_20220523_104016')
imgs = io.imread(image_cp)
channels = [[0,0]]
masks, flows, styles = model.eval(imgs, diameter=None, channels=channels)
io.save_to_png(imgs, masks, flows, image_cp)
```

[ ]:
```
image_path = f"{frame_path}/frame.tif"
image = ij.io().open(image_path)
imp = ij.py.to_imageplus(image)
input_txt = Path(f"{frame_path}/frame_cp_outlines.txt")
txt_fh = open(input_txt, 'r')
for line in txt_fh:
    xy = line.rstrip().split(",")
    xy_coords = [int(element) for element in xy if element not in '']
    x_coords = [int(element) for element in xy[::2] if element not in '']
    y_coords = [int(element) for element in xy[1::2] if element not in '']
    xcoords_jint = JArray(JInt)(x_coords)
    ycoords_jint = JArray(JInt)(y_coords)
    polygon_roi_instance = scyjava.jimport('ij.gui.PolygonRoi')
    roi_instance = scyjava.jimport('ij.gui.Roi')
    imported_polygon = polygon_roi_instance(xcoords_jint, ycoords_jint,␣
↪len(x_coords), int(roi_instance.POLYGON))
    imp.setRoi(imported_polygon)
    rm.addRoi(imported_polygon)
ij.py.run_macro("roiManager('Select All');")
```

```
rm.runCommand("Save", f"{frame_path}/" + f"frame.zip")
```

### 7 Display the segmentation result and selection of dead for signal cleaning

[ ]:
```
# show the result of the segmentation
imp.show()
ij.IJ.resetMinAndMax(imp)
rm.runCommand("Show All with labels")
ij.IJ.run("Brightness/Contrast...")
```

[ ]:
```
# enter label of the dead cells between the brackets
dead_cells = [1, 9]
```

[ ]:
```
# close all the images and clear the ROI manager
rm.runCommand("Select All")
rm.runCommand("Delete")
ij.py.run_macro('close("*")')
```

[ ]:
```
# open the isolated bacteria channel and clear the signal outside the cells and␣
↪inside the dead cells
image_path = f"{frame_path}/channel_bact.tif"
imp = ij.IJ.openImage(image_path)
imp.show()
input_ROI = f"{frame_path}/frame.zip"
rm.open(input_ROI)
rm.runCommand("Select All")
rm.runCommand("XOR")
ij.IJ.run("Clear Outside", "stack")
ij.IJ.run("Select None")
ij.IJ.run("Smooth", "stack")
ij.IJ.run("Smooth", "stack")
for cells in dead_cells:
    rm.select(cells-1)
    ij.IJ.run("Clear", "stack")
rm.runCommand("Select All")
rm.runCommand("Delete")
ij.IJ.run("Select None")
```

### 8 MCV centroid detection with Trackmate

[ ]:
```
import sys
import os

# Creation of output directory
```

```python
out = frame_path+"Output/"
if not os.path.exists(out):
    os.makedirs(out)

# Parameters for Detection
# Here, the user can specify parameters for the detection step in Trackmate␣
↪(Threshold Detector)
dsettings = {
    'TARGET_CHANNEL' : ij.py.to_java(2),
    'SIMPLIFY_CONTOURS' : False,
    'INTENSITY_THRESHOLD' : 5.0,
}

# Parameters for Tracking
# Here, the user can specify parameters for the tracking step in Trackmate (LAP␣
↪Tracker)
frame_gap = 2
tsettings = {
    'LINKING_MAX_DISTANCE' : 40.0,
    'ALLOW_GAP_CLOSING' : True,
    'GAP_CLOSING_MAX_DISTANCE' : 40.0,
    'MAX_FRAME_GAP' : ij.py.to_java(2),
    'ALLOW_TRACK_SPLITTING' : False,
    'SPLITTING_MAX_DISTANCE' : 15.0,
    'ALLOW_TRACK_MERGING' : False,
}

# Create Model
model = Model()
settings = Settings(imp)

# Detector
settings.detectorFactory = ThresholdDetectorFactory()
for parameter, value in dsettings.items():
    #settings.detectorSettings.update({parameter:value})
    settings.detectorSettings[parameter] = value
filter1 = FeatureFilter('QUALITY', 45, True)
settings.addSpotFilter(filter1)
print(settings.detectorSettings)

# Tracker
settings.trackerFactory = SparseLAPTrackerFactory()
settings.trackerSettings = settings.trackerFactory.getDefaultSettings()
for parameter, value in tsettings.items():
    #settings.trackerSettings.update({parameter:value})
    settings.trackerSettings[parameter] = value
```

```
# Execute Tracking
trackmate = TrackMate(model, settings)
ok = trackmate.checkInput()
if not ok:
    sys.exit(str(trackmate.getErrorMessage()))
ok = trackmate.process()
if not ok:
    sys.exit(str(trackmate.getErrorMessage()))
selectionModel = SelectionModel(model)

    # Display
ds = DisplaySettingsIO.readUserDefault()
#displayer = HyperStackDisplayer(model, selectionModel, imp, ds)
#displayer.render()
#displayer.refresh()
#trackscheme = TrackScheme(model, selectionModel, ds)
#trackscheme.render()

# Save Data
outFile = Path(out+"bact_exportModel.xml")
writer = TmXmlWriter(outFile)
writer.appendModel(model)
writer.appendSettings(settings)
writer.writeToFile()
csvFileTracks = Path(out+"bact_exportTracks.csv")
csvFileSpots = out+"bact_exportspots.csv"
#trackTableView = TrackTableView(model, selectionModel, ds)
#trackTableView.getTrackTable().exportToCsv(csvFileTracks)
#trackTableView.getSpotTable().exportToCsv(csvFileSpots)

spotsTableView = AllSpotsTableView(model, selectionModel, ds)
spotsTableView.exportToCsv(csvFileSpots)
```

### 9 ROI collection with Fiji and measurement of centroid coordinates

[ ]:
```
roi_collection = """
setAutoThreshold("Default dark no-reset");
run("Threshold...");
setThreshold(5, 255);
setOption("BlackBackground", true);
run("Convert to Mask", "black");
run("Analyze Particles...", "size=10-Infinity add stack");
run("Set Measurements...", "area centroid stack redirect=None decimal=2");
nbArea=roiManager("count")
for (i=0; i<nbArea; i++) {
```

```
        roiManager("Select", i);
        run("Measure");
}
close();
//close();
"""
rois = ij.py.run_macro(roi_collection)
f_name = os.path.basename(image_path)
f_name = os.path.splitext(f_name)[0]
rm.runCommand("Select All")
rm.runCommand("Save", f"{frame_path}/" + f"{f_name}.zip") # this saves the ROIs␣
↪as a zip file
rm.runCommand("Delete")
measurements = ij.ResultsTable.getResultsTable() # call of the table
measurements_table = ij.convert().convert(measurements, scyjava.jimport('org.
↪scijava.table.Table')) # conversion to a java table object
table = ij.py.from_java(measurements_table) # Conversion into a python␣
↪dataframe from Java
output_path = Path(f"{frame_path}/{f_name}.csv") # save giving a name matching␣
↪the opened image
table.to_csv(output_path)
ij.py.run_macro("""
    title = Table.title();
    selectWindow(title);
    run("Close");
""")
```

### 10 Matching of Trackmate spots results with with particles detection result

[ ]:
```
# Function for distance calculation between Trackmate spots centroids and Fiji␣
↪ROIs centroids
from math import isnan
def xref_locations(first, second, first_x='POSITION_X', first_y='POSITION_Y',␣
↪first_z='POSITION_Z',
                   second_x='X', second_y='Y', second_z='Slice',
                   max_dist=20, verbose=False):
    pairwise_elements = pandas.DataFrame()
    first_measurements = pandas.read_csv(first)
    first_measurements = first_measurements.drop([0,1,2])
    second_measurements = pandas.read_csv(second)
    first_gdf = geopandas.GeoDataFrame(
        first_measurements,
        geometry=geopandas.points_from_xy(first_measurements[first_x],
                                          first_measurements[first_y],
```

```
                                                first_measurements[first_z]))
    second_gdf = geopandas.GeoDataFrame(
        second_measurements,
        geometry=geopandas.points_from_xy(second_measurements[second_x],
                                          second_measurements[second_y],
                                          second_measurements[second_z]))

    ti_rows = first_gdf.shape[0]
    tj_rows = second_gdf.shape[0]
    for ti_row in range(0, ti_rows):
        if verbose:
            print(f"On row: {ti_row}")
        ti_element = first_gdf.iloc[[ti_row, ]]

        titj = geopandas.sjoin_nearest(ti_element, second_gdf,
                                       distance_col="pairwise_dist",
                                       max_distance=max_dist)
        chosen_closest_dist = titj.pairwise_dist.min()
        if (isnan(chosen_closest_dist)):
            print(f"This element has no neighbor within {max_dist}.")
        else:
            chosen_closest_cell = titj.pairwise_dist == chosen_closest_dist
            chosen_closest_row = titj[chosen_closest_cell]
            pairwise_tmp = pandas.concat([pairwise_elements,␣
↪chosen_closest_row])
            pairwise_elements = pairwise_tmp
    return pairwise_elements
```

[ ]:
```
#Running the function
import matplotlib.pyplot as plt
first = Path(f"{directory_path}/frameOutput/bact_exportspots.csv")
second = Path(f"{frame_path}/channel_bact.csv")
pairwise = xref_locations(first, second,
                          first_x='POSITION_X',
                          first_y='POSITION_Y',
                          first_z='POSITION_Z',
                          second_x='X',
                          second_y='Y',
                          second_z='Slice',
                          verbose=False)
#pairwise.head()
grouped = pairwise.groupby('ID')['Unnamed: 0'].apply(list).reset_index()
grouped.rename(columns={'Unnamed: 0': 'object_ID_list'}, inplace=True)
final_csv = Path(f"{frame_path}/channel_bact_grouped.csv")
grouped.to_csv(final_csv)
```

### 11 quality control of ROIs retained by nearest neighbor calculation

[ ]:
```
# This is to make the number of each lane of the "_grouped csv file" to be␣
↪usable for the ROI manager later
grouped_path = Path(f"{frame_path}/channel_bact_grouped.csv")
df = pandas.read_csv(grouped_path)
ROI_ID = []
for i in range(len(df)):
    roi = df['object_ID_list'][i]
    nums = roi.strip('[').strip(']')
    ROI_ID.append(int(nums))
print(ROI_ID)
print(len(ROI_ID))
```

[ ]:
```
# opening of the bacterial channel and the Fiji ROIs
bact_channel = f"{frame_path}/channel_bact.tif"
image = ij.io().open(bact_channel)
shown = ij.ui().show(image)
roi_input = f"{frame_path}/channel_bact.zip"
rm.open(roi_input)
```

[ ]:
```
# For each line of the grouped CSV file, a color is randomly picked and the␣
↪ROIs of the lines are overlayed on the image with the color.
import random
colors = ["blue", "cyan", "green", "magenta", "orange", "red", "white",␣
↪"yellow"]
for i in ROI_ID:
    random_color = random.choice(colors)
    rm.select(i)
    overlay_command = f"Overlay.addSelection('{random_color}', 2);"
    ij.py.run_macro(overlay_command)
```

[ ]:
```
# when done checking the quality of the tracking, this will close everything␣
↪and clears the ROI manager
rm.runCommand("Select All")
rm.runCommand("Delete")
ij.py.run_macro('close("*")')
```

#### 12 Fluorescence measurement on bacteria ROI in the other channels

[ ]:
```
# get the files list
file_pattern = os.path.join(directory_path, "*.tif")
file_list = glob.glob(file_pattern)

#set measurement
set_string = f'Set Measurements...'
measure_string = f'mean stack redirect=None decimal=2'
ij.IJ.run(set_string, measure_string)
cois = ['LC3', 'LV']

# preparation of variables for file name call automation
for file_path in file_list:
    f_name = os.path.basename(file_path)
    basename = os.path.splitext(f_name)[0]

    if basename.startswith(cois[0]):
        corename = basename.split("_", maxsplit=1)[1]
        input_roi = Path(f"{frame_path}/channel_bact.zip")
        rm.open(f"{input_roi}")

        for channel in cois:
            c_path = Path(f"{directory_path}/{channel}_{corename}.tif").
↪as_posix()
            image_c = ij.io().open(c_path)
            ij.ui().show(image_c)
            for i in ROI_ID:
                rm.select(i)
                ij.IJ.run('Measure')
            output = Path(f"{directory_path}/measurement/{channel}_{corename}.
↪csv").as_posix()
            saving = ij.IJ.saveAs("Results", output)
            ij.IJ.run("Clear Results")
            ij.py.run_macro('close("*")')
rm.runCommand("Select All")
rm.runCommand("Delete")
```

[ ]:
```
# this whole block is to merge the CSV files from the measurement part

cell_list = []
for filename in os.listdir(directory_path+"/measurement"):
    if filename.startswith(cois[0]):
        basename = filename[len(cois[0])+1:]
        corename = os.path.splitext(basename)[0]
        cell_list.append(corename)
```

```
for cell in cell_list:
    input_csv_lc3 = Path(f"{directory_path}/measurement/LC3_{cell}.csv")
    input_csv_lv = Path(f"{directory_path}/measurement/LV_{cell}.csv")

    #first marker csv
    df1 = pandas.read_csv(input_csv_lc3)
    df1.rename(columns={'Mean': 'Mean_LC3'}, inplace=True)
    df1 = df1.drop(columns = [df1.columns[0]], axis =1)

    #second marker csv
    df2 = pandas.read_csv(input_csv_lv)
    df2.rename(columns={'Mean': 'Mean_LV'}, inplace=True)
    df2 = df2.drop(columns =[df2.columns[0],df2.columns[2]], axis =1)

    #create new merged dataframe:
    final_results = pandas.concat([df1, df2], axis = 1)
    final_results = final_results.iloc[:, [1, 0, 2]]

    out = directory_path +"/final_output/"
    if not os.path.exists(out):
        os.makedirs(out)
    output_path = Path(f"{out}/{cell}_final_result.csv")
    final_results.to_csv(output_path)
```
